# Cell atlas of the human ocular anterior segment: Tissue-specific and shared cell types

**DOI:** 10.1101/2022.01.19.476971

**Authors:** Tavé van Zyl, Wenjun Yan, Alexi McAdams, Aboozar Monavarfeshani, Gregory S. Hageman, Joshua R. Sanes

**Author notes:** Correspondence to: Joshua R. Sanes or Tavé van Zyl. Equal contribution (Tavé van Zyl, Wenjun Yan). T.V.Z, Department of Ophthalmology and Visual Science, Yale School of Medicine, New Haven, CT. A.M.M., California Institute of Technology, Pasadena, CA 91125.

## Abstract

The anterior segment of the eye consists of the cornea, iris, ciliary body, crystalline lens and aqueous humor outflow pathways. Together, these tissues are essential for the proper functioning of the eye. Disorders of vision have been ascribed to defects in all of them; some, including glaucoma and cataract, are among the most prevalent causes of blindness in the world. To characterize the cell types that comprise these tissues, we generated an anterior segment cell atlas of the human eye using high throughput single-nucleus RNA sequencing (snRNAseq). We profiled 191,992 nuclei from non-diseased anterior segment tissues from 6 human donors, identifying >60 cell types. Many of these cell types were discrete, whereas others, especially in lens and cornea, formed continua corresponding to known developmental transitions that persist in adulthood. Having profiled each tissue separately, we performed an integrated analysis of the entire anterior segment revealing that some cell types are unique to single structure whereas others are shared across tissues. The integrated cell atlas was then used to investigate cell type-specific expression patterns of more than 900 human ocular disease genes identified either through Mendelian inheritance patterns or genome-wide association studies (GWAS).

**SIGNIFICANCE STATEMENT:** Several of the most prevalent blinding ocular conditions worldwide, including glaucoma, cataract and uncorrected refractive error, involve structures of the anterior segment of the human eye, which consists of the cornea, iris, ciliary body, crystalline lens and aqueous humor outflow pathways. In addition to providing transcriptomic profiles of the cell types within individual tissues, this work contributes to our understanding of the relatedness and diversity of these cell types across contiguous tissues by generating an integrated anterior segment cell atlas and documenting the expression of over 900 disease-associated genes in each cell type. By allowing simultaneous interrogation of cell-type specific expression of genes across multiple tissues, the atlas may yield broad insight into normal and disease-associated anterior segment functions.

## INTRODUCTION

The anterior segment of the eye is a complex set of interconnected structures, comprising the cornea, conjunctiva, iris, ciliary body, crystalline lens and aqueous humor outflow pathways; the outflow pathways, in turn, include the trabecular meshwork (TM), Schlemm canal and ciliary muscle (Fig. 1A, B). Together, these structures fulfill two prerequisites of vision: (1) ensuring that light reaches the retina, and (2) ensuring that it is optimally focused. The transparent cornea and lens provide the refractive power of the eye. The cornea also serves a critical barrier function. The iris determines how much light reaches the retina. The ciliary body produces a transparent fluid known as aqueous humor that nourishes the avascular cornea and lens and removes waste products. Finally, the outflow pathways drain aqueous humor from the anterior chamber at an appropriate rate to avoid abnormal elevations of intraocular pressure (IOP).

**Figure 1.**
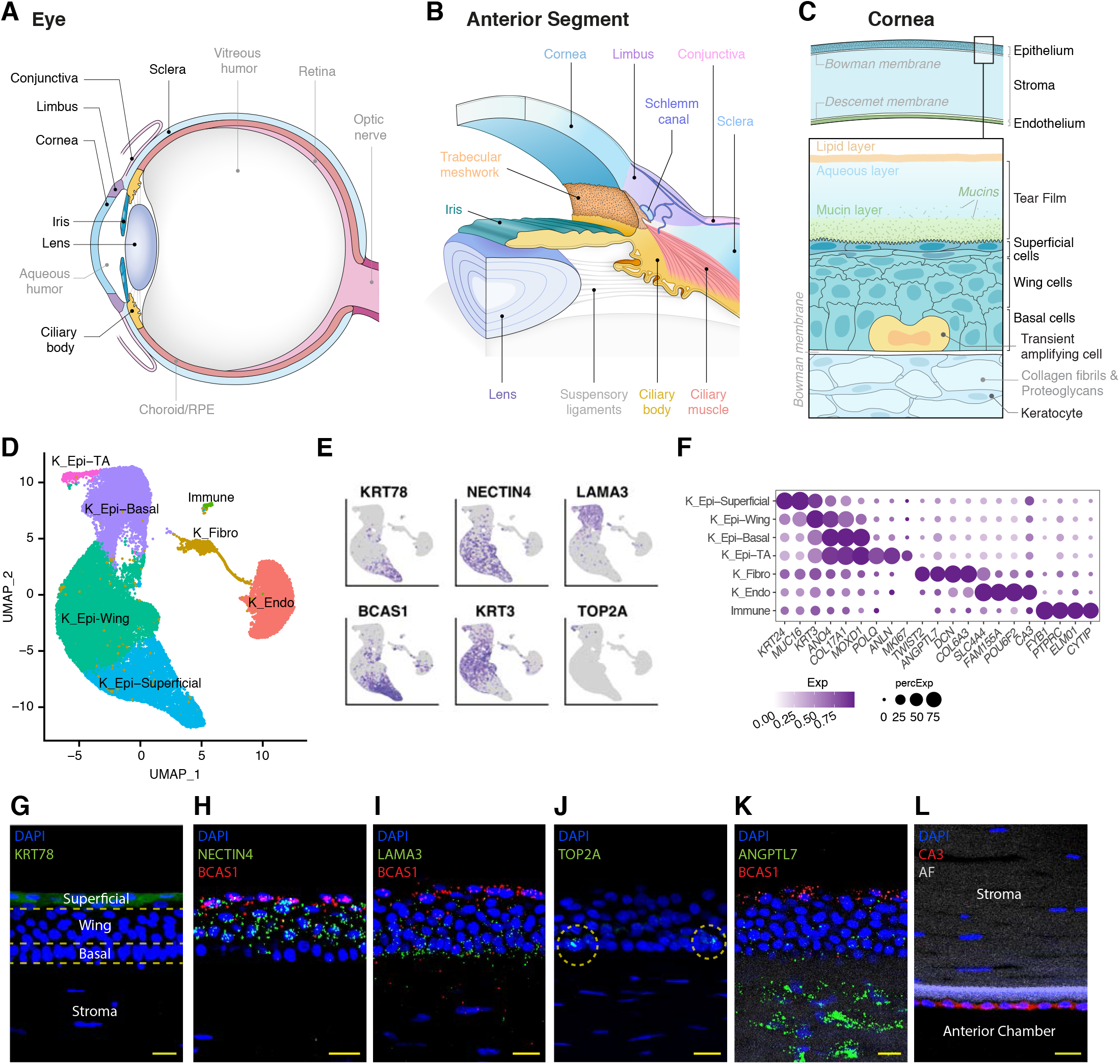
Human anterior segment and cells of the central cornea. **A** Diagram of the human eye, depicted in sagittal cross section. **B** Diagram of the anterior segment, which includes the cornea, iris, ciliary body and lens. The limbus, representing the transition between peripheral cornea and sclera, houses the aqueous outflow structures including the trabecular meshwork and Schlemm canal. **C** Diagram of the central cornea, comprised of 3 primary cellular layers: epithelium, stroma and endothelium. The multilayered epithelium is composed of basal, wing, and superficial cells delineated in boxed area. Transit amplifying cells are mitotically active basal cells. Keratocytes, specialized corneal fibroblast-like cells, are the main cell type within the stroma. **D** Clustering of 37,485 single-nucleus expression profiles from human central cornea visualized by Uniform Manifold Approximation and Projection (UMAP). Here and in subsequent UMAPs, arbitrary colors are used to distinguish clusters deemed to be distinct by unsupervised analysis. **E** Feature plots demonstrating representative differentially expressed (DE) genes corresponding to the epithelial subtypes. While superficial and basal cells had more discrete expression profiles, wing cells showed gradient expression reflecting their intermediate state. **F** Dot plot showing genes selectively expressed in cells of the central cornea, with gradient expression patterns noted in the epithelial subtypes. The small immune cell cluster likely represents a mixture of macrophages (or Dendritic cells) and lymphocytes. In this and subsequent figures, the size of each circle is proportional to the percentage of nuclei within a cluster expressing the gene and the color intensity depicts the average normalized transcript count in expressing cells. **G** Corneal superficial epithelium immunostained for KRT78 (green). **H** Fluorescent RNA *in situ* hybridization of *BCAS1* (red) highlights superficial epithelium and *NECTIN4* (green) highlights both wing cells and superficial cells. **I** Fluorescent RNA *in situ* hybridization of *BCAS1* (red) highlights superficial epithelium and *LAMA3* (green) highlights basal cells. **J** Transit Amplifying cells are identified via fluorescent RNA *in situ* hybridization for *TOP2A* (green). **K** Corneal stromal fibroblasts are highlighted by fluorescent RNA *in situ* hybridization for *ANGPTL7* (green); superficial epithelium highlighted by Fluorescent RNA *in situ* hybridization for *BCAS1* (red). **L** Corneal endothelium immunostained for CA3 (red). AF, autofluorescence; K_Epi, corneal epithelium; TA, transit amplifying; K_Fibro, corneal fibroblasts; K_Endo, corneal endothelium; DAPI, 4′,6-diamidino-2-phenylindole. Yellow bars show 20µm.

Because each of the structures within the anterior segment is essential for proper functioning of the eye, dysfunction of any one of them leads to vision loss. Indeed, the three leading causes of global blindness among adults ages >50 years involve anterior segment structures to varying extents: cataract (lens), glaucoma (outflow pathways), and uncorrected refractive error (lens and cornea) (Adelson 2020). Other important ophthalmologic conditions with manifestations primarily in the anterior segment include dry eye disease, corneal dystrophies, anterior uveitis, trachoma and onchocerciasis (river blindness). While most of these entities can be treated, few if any can be cured with current approaches.

To build a better understanding of the complex tissues comprising the human anterior segment, we generated a cell atlas using high throughput single-nucleus RNA sequencing (snRNAseq). We profiled 191,992 single nuclei from non-diseased anterior segment tissues, applied computational methods to cluster them based on transcriptomic similarity, and used histological techniques to assign cell type identities to the clusters. After investigating each dataset independently, we pooled them and performed an integrated analysis. In this way, we were able to show that some cell types are confined to specific tissues whereas others are shared across tissues. Finally, we used this cell atlas to investigate cell type-specific expression patterns of over 900 genes that have been implicated in susceptibility to human ocular diseases through Mendelian inheritance patterns or genome-wide association studies (GWAS).

## RESULTS

Six tissues – central cornea, corneoscleral wedge (CSW), trabecular meshwork, iris, ciliary body and lens – were dissected from eyes of six individual donors with no histories of ocular disease (*SI Appendix* Table S1). Tissues were obtained post-mortem within <6 hours from death in all but one case, dissected within an hour of enucleation, and frozen for further processing. Nuclei were prepared and profiled using a droplet-based method (Zheng, 2017). Altogether we obtained high-quality transcriptomes from 191,992 single nuclei from which we generated a cell atlas. An additional eight eyes were used for histological analysis (*SI Appendix* Table S2).

### Cornea

The transparent, avascular cornea forms a tough protective envelope, contiguous with the sclera, that encases the delicate intraocular tissues. Beyond its protective function as the first line of defense against infection and trauma, the cornea provides the principal refracting surface of the eye (Delmonte 2011). The cornea is composed of 3 primary cellular layers: epithelium, stroma, and endothelium (Fig. 1C). The corneal epithelium is a nonkeratinizing stratified squamous epithelium. Its 5-6 layers of cells can be divided into 3 histologically distinct sublayers: (1) an innermost single layer of columnar basal cells; (2) a 2-3 cell-thick intermediate layer of “wing” or polygonal suprabasal cells; and (3) a 2-cell thick superficial layer of plate-like squamous cells. The basal cell layer contains two cell types: some divide continuously (transit amplifying cells) while others arrest and subsequently migrate superficially to become wing and then surface cells. The avascular and acellular Bowman’s layer separates the epithelium from the lamellar stroma; and Descemet membrane, a basement membrane, separates the lamellar stroma from the endothelium.

We isolated the central 6 mm of the cornea and processed it separately from the peripheral cornea and limbus, which are discussed below. We recovered high quality transcriptomes from 37,485 single nuclei and divided them into 7 clusters using computational analysis (see Methods). Using established markers from the literature, we annotated them as four groups of corneal epithelial cells (81.3%), corneal endothelium (14.5%), stromal keratocytes (3.6%), and immune cells (0.5%) (Fig. 1D-F, *SI Appendix*, Fig. S1A).

To assign clusters to cell types, we localized differentially expressed genes by immunohistochemistry and *in situ* hybridization. We thereby demonstrated that the epithelial clusters corresponded to cells populating distinct epithelial sublayers. The basal epithelial cluster was characterized by enriched expression of ECM- and adhesion-related genes including *LAMA3* (Fig. 1E, I). The transit amplifying cells were distinguished from committed basal cells by selective expression of DNA repair and cell proliferation-associated genes including *TOP2A* (Fig. 1E, J) and *MKI67*. The superficial-most squamous epithelial cells were characterized by expression of multiple keratins (*KRT4, KRT19, KRT24, KRT78*) and mucins (*MUC4, MUC16, MUC21, MUC22*), as well as *BCAS1* (Fig. 1E, H, I, K). The middle layer wing cells were characterized by enriched expression of *KRT3* and graded expression of multiple genes, representing the transitional state between basal and superficial cells. Among them, *COL17A1, ANO4, KRT12, KC6* were expressed robustly in basal cells and weakly in the surface cells whereas *NECTIN4* and *MUC16* were absent from basal cells but expressed robustly by surface cells (Fig. 1E, F, H).

We also confirmed that two other clusters corresponded to corneal endothelium (*CA3;* Fig. 1F, L) and corneal stromal keratocytes (*ANGPTL7*; Fig. 1F, K). Finally, the small cluster of immune cells contained both macrophages (likely corneal dendritic cells) and lymphocytes; differentially expressed genes within the cluster included *PTPRC*/*CD45, CYTIP, FYB1, ELMO* and lymphocyte-associated marker *IKZF1* (Fig. 1F).

### Limbus

The limbus is the annular region between the avascular clear cornea and the vascularized opaque sclera (Fig. 2A). The limbus is not anatomically discrete but contains components distinct from both cornea and sclera (van Buskirk, 1989). Externally, it contains the transition zone of the ocular surface where corneal and conjunctival epithelia meet. Herein, limbal epithelial stem cells (LESC), accounting for less than 1-2% of proliferating basal epithelial cells, help maintain corneal epithelial integrity, which keeps the cornea avascular and transparent (Seyed-Safi, 2020; Lavker, 2020). Internally, the limbus includes the trabecular meshwork, Schlemm canal, and specialized outflow vessels that allow egress of aqueous humor from the anterior chamber.

**Figure 2.**
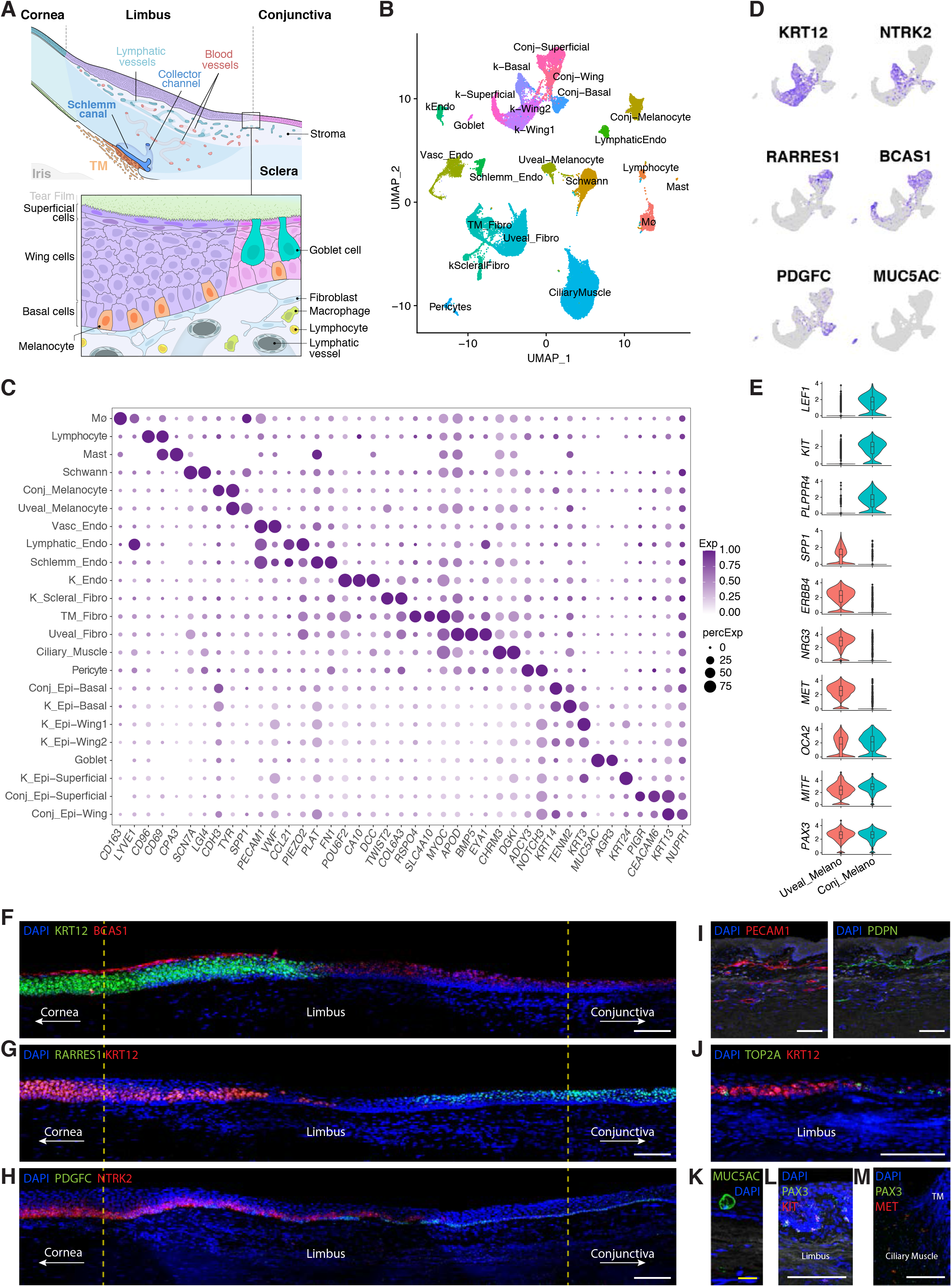
Cell types of the internal and external limbus derived from dissection of corneoscleral wedge (CSW) **A** Diagram of the limbus, representing the transitional tissue between peripheral corneal and sclera. Externally, the corneal epithelium transitions into conjunctival epithelium with a vascularized subepithelial stroma, delineated in boxed area. Internally, the aqueous humor outflow structures, including trabecular meshwork, Schlemm canal and the collector channels are noted. **B** Clustering of 51,306 single-nucleus expression profiles derived from corneoscleral wedge tissue visualized by Uniform Manifold Approximation and Projection (UMAP). **C** Dot plot showing genes selectively expressed in cells of the internal and external limbus. **D** Feature plots showing a selection of genes enriched in ocular surface epithelium subtypes. **E** Violin plot showing common and differentially expressed genes expressed in conjunctival and uveal melanocytes. **F** Fluorescent RNA *in situ* hybridization for *BCAS1* (red) highlights superficial epithelium in both the cornea and conjunctiva while *KRT12* (green) highlights basal and wing cells of the cornea. A transitional area is noted in the limbus where *KRT12* expression tapers off and is absent in the conjunctival epithelium. **G** Fluorescent RNA *in situ* hybridization for *KRT12* (green) highlights basal and wing cells of the cornea and *RARRES1* (green) highlights superficial and some wing cells of the conjunctiva. A transition area within the limbus is noted. **H** Fluorescent RNA *in situ* hybridization for *NTRK2* (red) highlights basal epithelium of the cornea and *PDGFC* (green) highlights mostly basal epithelium of the conjunctiva. **I** Immunostaining against PECAM1 (red) highlights endothelial cells lining vessels in the subepithelial stromal tissues of the external limbus. Immunostaining against PDPN (green) highlights lymphatic endothelium lining a subset of these vessels. **J** Transitly amplifying basal cells within the limbus are highlighted with Fluorescent RNA *in situ* hybridization for *TOP2A* (green); basal and wing cells in the limbal area demonstrate KRT12 (red) expression as visualized by Fluorescent RNA *in situ* hybridization. **K** A goblet cell in the conjunctiva visualized with immunostaining against MUC5AC (green). **L** Conjunctival melanocytes visualized by fluorescent RNA *in situ* hybridization for PAX3 (green) and KIT (red). **M** Uveal melanocytes visualized by fluorescent RNA *in situ* hybridization for PAX3 (green) and MET (red). Mø, macrophage; Conj, conjunctival; Vasc, vascular; Endo, endothelium, K_Endo, corneal endothelium; K_Scleral_Fibro, corneoscleral fibroblast; TM, Trabecular Meshwork; Fibro, fibroblast; Conj_Epi, conjunctival epithelium; K_Epi, corneal epithelium; DAPI, 4′,6-diamidino-2-phenylindole. Scale bars show 100µm.

To ensure adequate representation of cell types in both the internal and external limbus, we separated the trabecular meshwork (TM) from the corneoscleral wedge (CSW) and processed each portion separately, then merged the data into a single dataset of 51,306 transcriptomes due to the large overlap of cell types (Fig. 2B, C; *SI Appendix*, Fig. S1B, S2F). Of these, 10,510 were identified as ocular surface epithelial cells based on expression of canonical epithelial markers *CDH1, CLDN1* and keratin genes; we re-clustered this group to identify rare types and ensure optimal capture of gradient differences (*SI Appendix*, Fig. S1C, S2A). This clustering result was then incorporated into the limbus dataset.

#### Ocular Surface Epithelium

We identified seven groups of ocular surface epithelial cells, which could be divided into corneal and conjunctival populations based on established markers: those with robust expression of *KRT12, KRT3* and/or *KRT24* were corneal, whereas those with *AQP3, AQP5* and/or *KRT13* expression were conjunctival (*SI Appendix*, Fig. S2B). The corneal clusters corresponded to the superficial, basal and wing types described above (Figure 1D), but in this case, two groups of wing cells were discernible (K_Epi-Wing1 and K_Epi-Wing2). K_Epi-Wing1 exhibited expression patterns similar to that of the corneal superficial epithelium and K_Epi-Wing2 shared markers with basal epithelium of the cornea and conjunctiva (*SI Appendix*, Fig. S2 A and B). The three conjunctival clusters were identifiable as basal (e.g., *KRT14*), superficial (e.g., *MUC4, CEACAM6*) and wing, with graded differences reflecting the transition from basal to superficial cells. Two putative additional cell types did not form discrete clusters. One, the basal transit amplifying cells, were captured (Fig. 2J, *SI Appendix*, Fig. S2D), but did not form a distinct cluster in this smaller dataset (2718 basal epithelia in limbus compared to 8614 in central cornea). Likewise, a small subset of putative LESCs or limbal progenitor cells (*GPHA2+, CDH19+, GJA1-*) formed a discrete lobe within in the conjunctival basal cluster (*SI Appendix*, Fig. S2E) (Collin 2021; Ligocki 2021; Chen 2006). Thus, all together, we document the existence of nine epithelial cell groups in the limbus.

Corneal and conjunctival epithelium were similar in many respects but differed in others. For example, superficial cells of both the cornea and conjunctiva expressed *BCAS1, KRT4* and *MUC4*; superficial and wing cells of both the cornea and conjunctiva expressed *NECTIN4*; and basal cells of both the cornea and conjunctiva expressed *LAMA3* (*SI Appendix*, Fig. S2B). Regarding differences, corneal basal and wing epithelial cells in the cornea were strongly positive for *KRT12*, whereas their corresponding types in the conjunctiva were *KRT12*-negative (Fig. 2D, F, G, J). Similarly, corneal basal epithelial cells were strongly *NTRK2*-positive, whereas those in the conjunctiva were *NTRK2-*negative but positive for *PDGFC* and *LGR6* (Fig. 2D,H, *SI Appendix*, Fig. S2C, E). *BCAS1* expression was limited to the superficial epithelial layer in the cornea, but present in both the superficial and wing epithelial cells of the conjunctiva (Fig. 2F). These layers of the conjunctiva were also positive for *RARRES1* and *MECOM*, unlike their counterparts in the cornea (Fig. 2G, *SI Appendix*, Fig. S2C).

#### Goblet Cells

A distinct cluster corresponding to the specialized, mucin-producing goblet cells of the ocular surface was identified based on enrichment of *MUC5AC*, encoding a gel-forming mucin (Gipson 2016) (Fig. 2K). This cluster was localized to the conjunctival epithelium and also expressed *ATP2C2, AGR3*, and *TFF1*.

#### Melanocytes

Two distinct clusters were composed of melanocytes. They shared canonical melanocyte markers such as *PAX3, TYR*, and *MITF*, but were markedly distinct at a transcriptional level (Fig. 2C, E). Histology supported the presence of two distinct cell types, which we annotate as uveal and conjunctival, corresponding to their locations (Fig 2L, M). The uveal type, derived from both the TM strip and CSW, represented melanocytes in the ciliary muscle and selectively expressed *ERBB4, NRG3* and *MET*; the conjunctival type, derived exclusively from the CSW, represented melanocytes nestled among the basal epithelial cells of the conjunctiva and selectively expressed *LEF1, PLPPR4* and *KIT*.

#### Trabecular Meshwork

Two clusters of closely related cells demonstrated differentially expressed genes previously identified via scRNAseq analysis of TM tissue, including *CHI3L1, ANGPTL7, RSPO4, TMEFF2, PPP1R1B, BMP5* and *C7* (van Zyl et al 2020, Patel et al 2020). *In situ* hybridization confirmed localization of these genes to the aqueous drainage structures within the iridocorneal angle, with some clearly confined to the TM and others extending into contiguous tissues of the iris root and ciliary muscle (*SI Appendix*, Fig. 2 H-J). The two clusters were tentatively annotated as TM Fibroblasts and Uveal Fibroblasts, with the latter characterized by genes most strongly expressed at the uveal base of the TM, the iris root and ciliary muscle. Gradient expression patterns were noted within and across both clusters (*SI Appendix*, Fig. S2G), potentially representing further specialization as described below (Section on Integrated Analysis).

#### Vessel Endothelium

Transcriptomic profiles identified three clusters of vessel endothelial cells. Based on our previous work (van Zyl et al., 2020), we identified one as Schlemm canal endothelium expressing *FN1* and *PLAT* and a second as vascular endothelium, expressing VWF (Fig. 2C). A distinct sub-cluster within this vascular endothelial cluster corresponded to collector channels/aqueous veins (expressing *ACKR1/DARC, AQP1, SELE*, and *COL15A1*; see van Zyl et al 2020). The third endothelial cluster selectively expressed lymphatic markers including *LYVE1, PROX1, CCL21* and *FLT4* and was localized to subepithelial vessels within the conjunctival stroma using immunostaining against PDPN (Fig. 2I). We assigned these cells to the conjunctival lymphatic endothelium.

#### Additional Cell Types

Based on marker expression, seven additional clusters were identified as pericytes (*NOTCH3* and *PDGFRB*), macrophages (*LYVE1*+*CD163*+), lymphocytes (*CD69*+), mast cells (*IL1RL1*+*CPA3*+), ciliary muscle cells (*DES* and *CHRM3*), Schwann cells (*LGI4* and *CDH19*), and a group of fibroblasts that likely included both corneal stromal fibroblasts (*KERA* and *MME*) and scleral fibroblasts (*TXNB, FBLN1*) (Fig. 2C).

### Iris

The iris, positioned along a plane that divides the anterior and posterior chambers, functions as a diaphragm akin to those in manufactured optical systems. The iris can be divided into five principal parts: (1) the anterior border layer, consisting of a dense meshwork of fibroblasts and melanocytes; (2) the stroma, made up of loose connective tissue and a lower density of fibroblasts and melanocytes, along with blood vessels, axons surrounded by Schwann cells, and immune cells such as macrophages (“clump cells”), mast cells and lymphocytes; (3) the sphincter muscle, made up of spindle-shaped (mononucleated) smooth muscle cells bundled into units of 5-8; (4) the dilator muscle, a group of pigmented myoepithelial cells sometimes called anterior pigmented epithelium; and (5) the posterior pigmented epithelium facing the posterior chamber (Fig. 3A).

**Figure 3.**
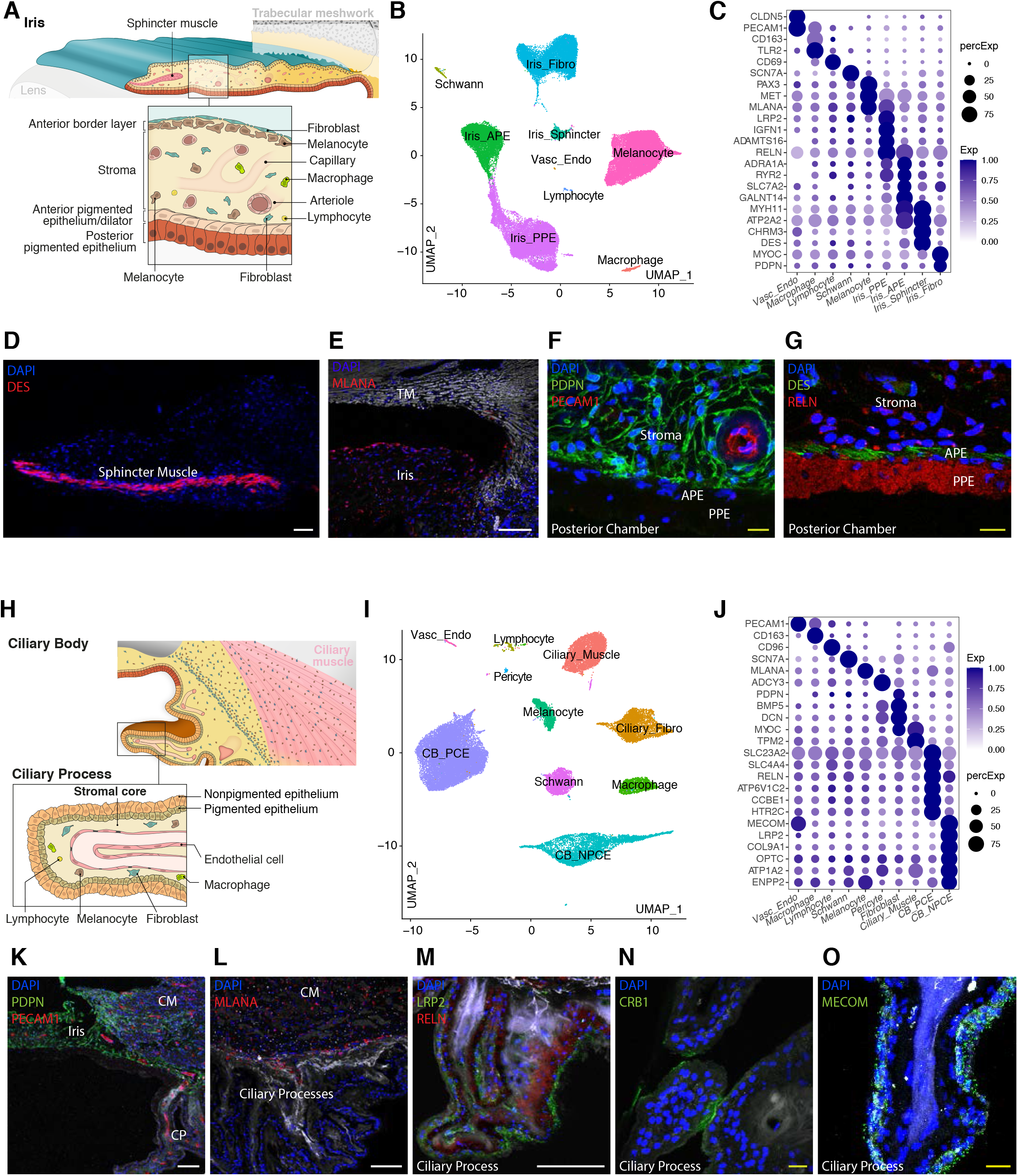
Cells of the iris and ciliary body. **A** Diagram of the human iris, consisting of the anterior border layer, the stroma, the sphincter muscle, anterior pigmented epithelium (APE) and posterior pigmented epithelium, shown in greater detail within box. **B** Clustering of 57,422 single-nucleus expression profiles derived from iris tissue visualized by Uniform Manifold Approximation and Projection (UMAP). **C** Dot plot showing genes selectively expressed in cells of the iris. **D** Iris sphincter muscle cells immunostained with DES (red). **E** Uveal melanocytes within the anterior border layer and stroma of the iris immunostained with MLANA (red). **F** Iris fibroblasts and vessel endothelium within the iris stroma immunostained with PDPN (green) and PECAM1 (red), respectively. **G** Immunostaining against RELN (red) highlights the iris posterior pigmented epithelium (PPE) and to a lesser extent iris anterior pigmented epithelium (APE); the latter is also positive for DES (green), highlighting its contractile role as the iris dilator. **H** Diagram of the human ciliary body, consisting of the ciliary muscle, ciliary stroma and ciliary processes, shown in greater detail within box. **I** Clustering of 34,132 single-nucleus expression profiles derived from ciliary body tissue visualized by Uniform Manifold Approximation and Projection (UMAP). **J** Dot plot showing genes selectively expressed in cells of the ciliary body. **K** Ciliary stromal fibroblasts immunostained with PDPN (green) and vessel endothelium with PECAM1 (red). **L** Uveal melanocytes within the ciliary stroma immunostained with MLANA (red). **M** Ciliary body non-pigmented ciliary epithelium (CB-NPCE) and pigmented ciliary epithelium (CB-PCE) immunostained with LRP2 (green) and RELN (red), respectively. **N** Immunostaining against CRB1 (green) highlights a subset of NPCE situated in proximity with neighboring ciliary processes. **O** RNA *in situ* hybridization for *MECOM* (green) highlights NPCE lining a ciliary process. Vasc_Endo, vascular endothelium; Fibro, fibroblast; CM, ciliary muscle, CP, Ciliary Process; DAPI, 4′,6-diamidino-2-phenylindole. White scale bars show 100µm; yellow bars show 20µm.

From iris samples, we obtained 57,422 nuclei that formed 9 clusters (Fig. 3B, C, *SI Appendix*, Fig. S1D). We were able to assign all 9 to cell types. Iris stromal fibroblasts, expressing *DCN, APOD, MYOC, RARRES1*, and *PDPN*, were identified in the stroma and anterior border layer of the iris via immunostaining against PDPN (Fig. 3F). The melanocytes were identified through their selective expression of melanocyte marker *PAX3*, as well as *EDNRB, MITF, MLANA, MLPH* and *TYR* (Fig. 3C,E). Smooth muscle cells comprising the iris sphincter muscle, responsible for constricting the pupil upon cholinergic stimulation, were identified by selective expression of the muscarinic receptor *CHRM3*, as well as *DES* (Fig. 3C,D). The iris dilator muscle, responsible for dilating the pupil upon adrenergic stimulation, selectively expressed the alpha-1a adrenergic receptor, *ADRA1A*. Both dilator and sphincter muscle cells expressed contractile genes, including *MYH11* and *ATP2A2* (Fig. 3C). The large, intensely pigmented cells of the posterior pigmented epithelium were visualized with immunostaining against RELN (Fig. 3G). Vascular endothelial cells, macrophages, lymphocytes and Schwann cells were identified by expression of canonical markers (see above and van Zyl et al., 2020).

### Ciliary Body

The ciliary body consists of four main components: (1) ciliary muscle, responsible for accommodation and assistance in removal of aqueous humor; (2) stroma, the vascularized connective tissue core; (3) pigmented ciliary epithelium (PCE); and (4) non-pigmented ciliary epithelium (NPCE). The PCE and NPCE are single-layered epithelia that lie adjacent to each other, oriented apex-to-apex and connected by gap junctions (Fig. 3H). Both play important roles in maintaining the blood-aqueous barrier and secreting aqueous humor.

Analysis of 34,132 ciliary body single nucleus transcriptomes yielded 10 distinct clusters (Fig. 3I, J, *SI Appendix* Fig. S1F). The most abundant corresponded to the PCE and NPCE. Differentially expressed genes in the PCE included *ATP6V1C2*, encoding the H^+^ vacuolar ATPase; *SLC23A2*, encoding a sodium-dependent ascorbate transporter; and *SLC4A4*, encoding the sodium-bicarbonate cotransporter (NBC) (Fig. 3J). Other differentially expressed genes included *MAMDC1, CCBE1, DCT* and numerous additional solute transporters (*SLC35G1, SLC38A11, SLC24A5, SLC9B2, SLC7A2, SLC7A6*), consistent with the PCE’s role alongside the NPCE in secreting aqueous humor. We visualized the PCE via immunostaining against RELN (Fig. 3M). Differentially expressed genes that distinguished NPCE from PCE included the vitreous humor components, *COL9A1, COL9A3* and *OPTC*, reflecting the NPCE’s close association with this gel (Le Goff, 2008), as well as *ATP1A2, NECTIN3, CACNA1E, ENPP2*, and the chloride transporter, *BEST2*. The NPCE was visualized with immunostaining against LRP2, MECOM and CRB1, the latter being expressed by a subset of cells populating areas of the ciliary processes which were in contact with other processes (Fig. 3M-O).

Cell types comprising the other clusters were determined by expression of canonical markers. They included *DES* and *MYH11* for muscle, *DCN* and *PDPN* for fibroblasts (Fig. 3K), *LGI4* and *SCN7A* for Schwann cells, *MLANA* for melanocytes (Fig. 3L), PECAM1 for vascular endothelium (Fig. 3K), *ADCY3* for pericytes, *CD163* for macrophages, and *CD96* for lymphocytes (Fig. 3J).

### Lens

The lens occupies the space between the iris and the vitreous (Fig. 1A). It is suspended in position by the elastic zonular fibers running between the equatorial lens capsule and the ciliary processes (Bassnett 2021) (Fig. 1B). The lens substance, composed of the central nucleus, concentrically layered cortical fibers, and lens epithelium, is enveloped by an elastic basement membrane called the lens capsule (Fig. 4A). Throughout life, lens epithelial cells that retain the capacity to divide migrate toward the post-equatorial region, where they terminally differentiate into cortical lens fiber cells (Bassnett & Sikic, 2017; Mochizuki & Masai, 2014). These are laid down in a compact lamellar fashion adjacent to previously formed fibers. Mature lens fiber cells lack most organelles, including the nucleus, ribosomes and mitochondria, rendering them highly transparent.

**Figure 4.**
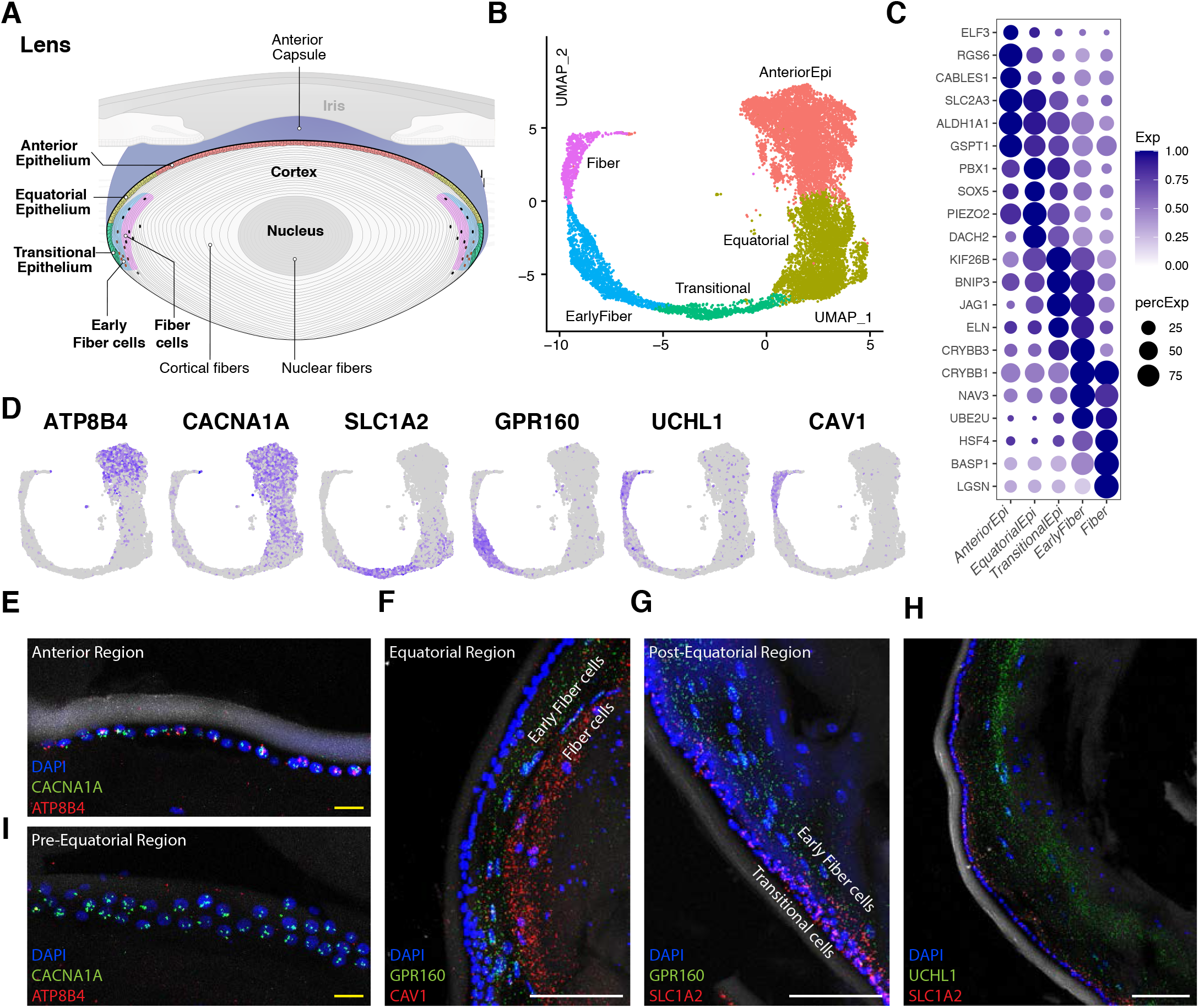
Cells of the crystalline lens. **A** Diagram of the human crystalline lens, consisting of lens epithelium and lens fiber cells. **B** Clustering of 13,900 single-nucleus expression profiles derived from lens tissue visualized by Uniform Manifold Approximation and Projection (UMAP). Rather than spatially discrete clusters, a continuum is observed ranging from anterior epithelium to lens fiber cells. **C** Dot plot showing genes selectively expressed in cells of the lens. **D** Feature plots demonstrating differentially expressed genes corresponding to cells of the lens. **E** Lens epithelial cells visualized with RNA *in situ* hybridization. Expression of *CACNA1A* (green) is evident in both anterior and equatorial epithelial cells whereas *ATP8B4* (red) is confined to the anterior epithelium. **F** RNA *in situ* hybridization demonstrates expression of *GPR160* (green) by early lens fiber cells, and *CAV1* (red) by more mature fiber cells. **G** Transitional lens epithelial cells in the post equatorial region are positive for *SLC1A2* (red) as visualized by RNA *in situ* hybridization; early fiber cells are positive for *GPR160* (green). **H** Transitional lens epithelial cells most abundant in the post equatorial region are positive for *SLC1A2* (red) as visualized by RNA *in situ* hybridization; fiber cells are positive for *UCHL1* (green). **I** Pre-equatorial lens epithelial cells are CACNA1A (green) positive but ATP8B4 (red) negative. White scale bars show 100µm; yellow bars show 20µm.

Computational analysis divided 13,900 lens cell transcriptomes into 5 clusters (Fig. 4B, *SI Appendix*, Fig. S1E). Three of the 5 clusters were classified as lens epithelial cells due to their robust expression of the ocular epithelial marker, *PAX6*. Their regional localizations as visualized by *in situ* hybridization for selectively expressed markers (*ATP8B4, CACNA1A1*, and *SLC1A2*) identified them as regionally enriched Anterior, Equatorial and Transitional types (Fig. 4C-E, G-I). Of note, cells in the transitional cluster expressed genes supporting differentiation into fiber cells (e.g., *JAG1, NAP1L4, BACH2*) (Le, 2009, Tanaka 2019, Zhao 2018). The remaining 2 clusters, visualized with *in situ* hybridization for *GPR160, CAV1* and *UCHL1*, corresponded to nucleated fiber cells at different stages of maturity (Fig. 4F-H).

The arrangement of these clusters strikingly mirrored the migratory and developmental trajectory of lens epithelial cells. Rather than being discrete, the clusters were continuous, describing a gradient leading from epithelial cell on one end of the spectrum to lens fiber cell on the other, as well as gradients of expression within each cluster (Fig. 4C and D).

### Integrated Analysis

In several cases, cell types identified in one tissue resembled those in one or more other tissues. This observation raised the question of which cell types were present in multiple tissues, and which types were tissue-specific. To address this issue, we pooled and reclustered data from iris, CB, cornea, CSW and TM. Initial analysis indicated that cell types in the lens were distinct from those in the other structures, so these were omitted from this analysis. Altogether, integration yielded 34 clusters (Fig. 5A, *SI Appendix*, Fig. S3A, S4A). A list of the differentially expressed (DE) genes from each type, as well as those from lens, is compiled in Supplementary Table 4.

**Figure 5.**
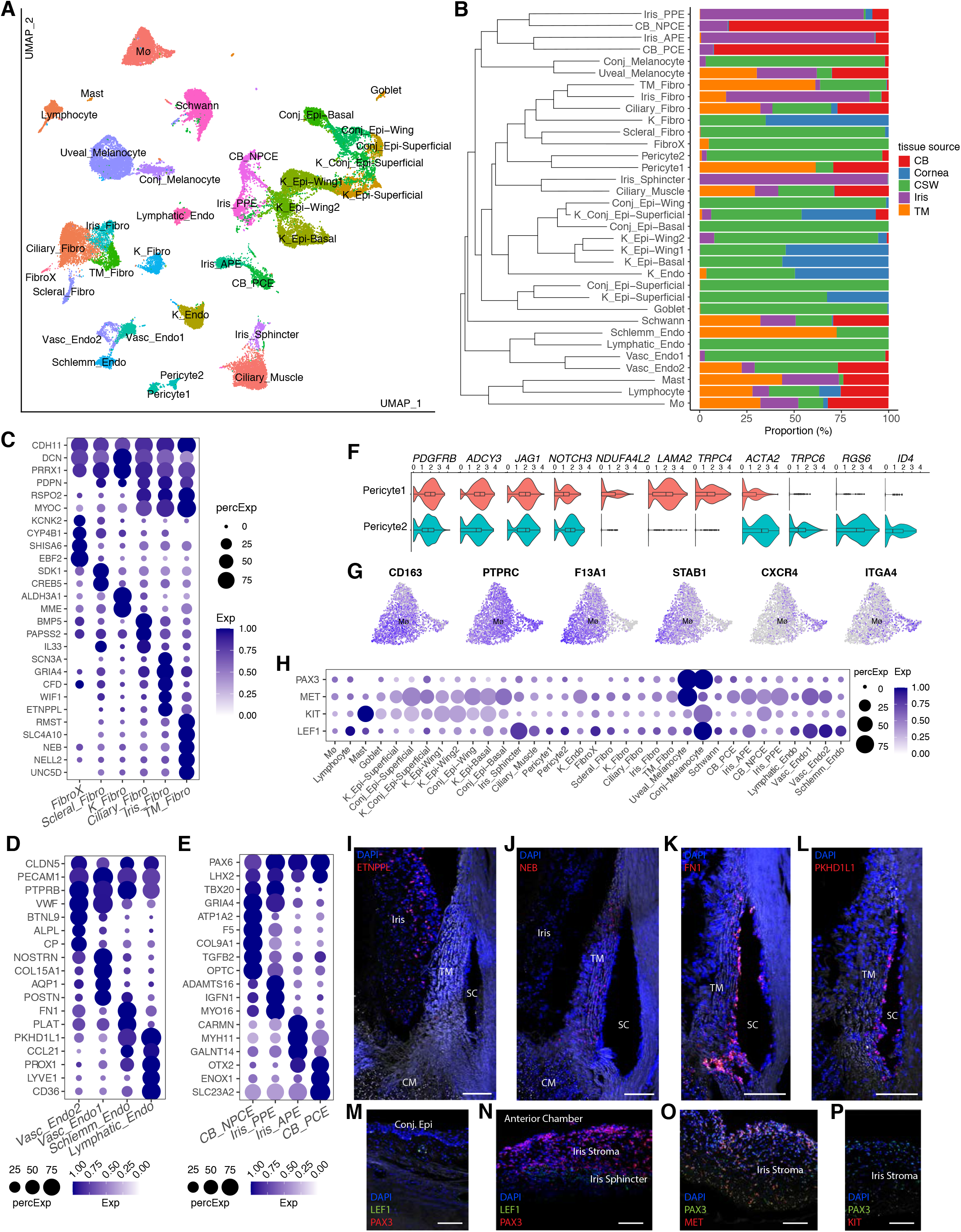
Integrated analysis of cells populating the human anterior segment. **A** Clustering of 42,901 single-nucleus expression profiles pooled together for the integrated analysis, visualized by Uniform Manifold Approximation and Projection (UMAP). **B** Stacked bar chart indicating proportions within each cluster contributed by separate tissue sources, including the cornea, corneoscleral wedge (CSW), iris and ciliary body. Transcriptional relatedness indicated by dendrogram. **C** Dot plot showing common and differentially expressed genes in fibroblast types. **D** Dot plot showing common and differentially expressed genes in vessel endothelial types. **E** Dot plot showing common and differentially expressed genes in uveal epithelium of the iris and ciliary body. **F** Violin plots showing common and selectively expressed genes in the two pericyte clusters. **G** Feature plots demonstrating expression patterns within the macrophage cluster. **H** Dot plot showing cell-type specific expression of PAX3, MET, KIT, and LEF1. **I** Iris fibroblasts selectively expressing *ETNPPL* (red), visualized with RNA *in situ* hybridization. **J** Trabecular meshwork (TM) fibroblasts selectively expressing *NEB* (red), visualized with RNA *in situ* hybridization. **K** Schlemm canal (SC) endothelium expressing FN1 (red), visualized with RNA *in situ* hybridization. **L** Schlemm canal (SC) endothelium expressing PKHD1L1 (red), visualized with RNA *in situ* hybridization. **M** Melanocytes within the basal layer of the conjunctiva visualized with RNA *in situ* hybridization for *PAX3* (red) and *LEF1* (green). Consistent with expression pattern seen in (H), vascular endothelium lining a vessel within the conjunctival stroma is also noted to be positive for *LEF1* (green). **N** Melanocytes within the iris stroma, visualized with RNA *in situ* hybridization for *PAX3* (red), do not express *LEF1* (green, absent). The iris sphincter is instead positive for *LEF1*, consistent with expression noted in (H). **O** Uveal melanocytes within the iris anterior border layer and stroma visualized with RNA *in situ* hybridization for *MET* (red) and *PAX3* (green). **P** Melanocytes within the iris stroma, visualized with RNA *in situ* hybridization for *PAX3* (red), do not express *KIT* (green, absent). Abbreviations as in previous figures. White scale bars show 100µm.

We then tallied the origins of the cells within each cluster. In many cases, cells of a particular type were derived mostly or entirely from a single tissue – for example PCE cells from the ciliary body, sphincter muscle from the iris, and goblet cells from the CSW (Fig. S3B and 5B). In some cases, however, cells derived from multiple tissues co-clustered. This was not surprising for immune cells, melanocytes and Schwann cells, but unexpected for some other cell types.

#### Epithelium

Twelve clusters were identified as epithelial cells. Most were tissue-specific, deriving primarily from iris, ciliary body, CSW, or cornea/CSW, with the latter group presumably reflecting the presence of corneal tissue in the CSW (Fig. 5B). Based on transcriptomic similarity they formed two clades. One comprised corneal and conjunctival types, and the other comprised uveal (iris and ciliary body) types, consistent with their embryological origins from the surface ectoderm or the neuroectoderm of the optic cup, respectively (Cvekl & Tamm, 2004). Transcriptional similarity further mirrored embryological origin among the four uveal epithelial types: the NPCE of the ciliary body was transcriptionally more similar to the posterior pigmented epithelium of the iris, together representing the inner layer of the uveal bilayer (derived from the inner layer of the optic cup), and the PCE of the ciliary body was more similar to the anterior pigmented epithelium of the iris, together representing the outer layer of the uveal bilayer (derived from the outer layer of the optic cup) (Fig. 5E).

#### Endothelium

Four clusters of vessel endothelial cells were identified through expression of the canonical marker *PECAM1* (Fig. 5D). All four clusters were transcriptomic neighbors, and three of the four were largely tissue-specific (Fig. 5B). Two corresponded to vascular endothelium, likely reflecting different portions of the vascular tree. One of these, Vasc_Endo1, was predominantly derived from the CSW tissue and expressed markers previously associated with collector channels and aqueous veins (*AQP1, POSTN, COL15A1*); it may also contain endothelium lining other vessels of the venous tree (van Zyl et al 2020). The other, Vasc_Endo2, was present in multiple tissues and may correspond to capillary endothelium and/or endothelium lining vessels of the arterial tree (Schupp et al, 2021). Two other clusters corresponded to lymphatic endothelium (*PDPN*+, *PKHD1L1*+, *SI Appendix*, Fig. S4G) predominantly localized to the conjunctival subepithelial stroma, and to Schlemm canal endothelium (*FN1+ PLAT+*, Fig. 5D, K). The predominantly TM source of this latter cluster supports the notion that Schlemm canal endothelium is a unique endothelium specialized for the conventional outflow path. Immunostaining against PKHD1L1 highlighted closely related conjunctival lymphatics and Schlemm canal endothelia (Fig. 5L) but not blood vessels.

#### Pericytes

Two distinct pericyte clusters were identified by shared expression of *PDGFRB, ADCY3, JAG1 and NOTCH3* (Fig. 5F). Their tissue origins were distinct: Pericyte1 (*LAMA2+*, TRPC4+), was derived mostly from TM and CB whereas Pericyte2, (*ID4+, TRPC6+, RGS6+*), was derived almost exclusively from the CSW (Fig. 5B *SI Appendix*, Fig. S3B, S4H-K).

#### Fibroblasts

Six transcriptomically related clusters expressed genes diagnostic of fibroblasts (e.g., *PDGFRA, COL6A3, DCN, PRRX1, CDH11;* Fig. 5B, C). Five of the clusters were assigned based on tissue source and histological validation: TM fibroblasts derived from tissue within the TM and CSW dissections; iris fibroblasts derived predominantly from the iris; Ciliary Fibroblasts derived primarily from the ciliary muscle tissue within the TM, CSW and CB (annotated as “uveal fibroblasts” in the TM/CSW atlas, above); corneal fibroblasts, derived predominantly from cornea and corneal component of the CSW dissection; and scleral fibroblasts derived exclusively from the scleral and/or limbal component of the CSW dissection. The sixth cluster, provisionally labeled Fibro X, was transcriptomically most similar to scleral fibroblasts and was derived almost exclusively from the CSW but exhibited distinct markers such as *EBF2* and *SHISA6*; this cluster remains incompletely characterized.

Three of the fibroblast clusters – iris, ciliary and TM fibroblasts – were closely related and shared numerous markers (e.g., *MYOC, PDPN*, and *RSPO2*). However, each expressed distinct markers that enabled histological validation to confirm their annotation based on tissue source: *WIF1* and *ETNPPL* for iris fibroblasts, *BMP5, PI16* and *C7* for ciliary fibroblasts, and *NEB, NELL2, UNC5D and TMEM178A* for trabecular meshwork fibroblasts (Fig. 5I,J, *SI Appendix*, Fig. S2I,J and S4C-F). Together, these clusters may be considered “outflow fibroblasts,” populating a key area within the iridocorneal angle where aqueous drains either through the conventional or uveoscleral pathway.

Surprisingly, unsupervised analysis did not further subdivide the TM cluster into beam and juxtacanalicular (JCT) cells as expected from studies using scRNAseq (van Zyl et al., 2020; Patel et al., 2020). However, heterogeneity was evident both within the TM cluster and across the iris and CB fibroblast clusters (*SI Appendix*, Fig. S4B). Markers of Beam A (e.g. BMP5) described in van Zyl et al. (2020) were expressed predominantly by the ciliary fibroblast cluster representing cells populating the ciliary muscle and uveal base of the TM; markers of Beam B (*TMEFF2*) and JCT (*CHI3L1*) were expressed by largely non-overlapping populations within the TM fibroblast cluster and exhibited correspondingly non-overlapping staining on histology (Fig. S3F, S2H-J). Together this suggests that the TM fibroblast cluster likely contains both beam and JCT cells and that the current dataset does not easily separate these two types.

#### Melanocytes

In the CSW atlas described earlier, two types of melanocytes were identified: conjunctival and uveal, with the latter deriving from the ciliary muscle component of the CSW dissection. Integrated analysis supported this distinction: melanocytes derived from the iris and ciliary body co-clustered with the uveal melanocytes (*MET*+) identified in the CSW, and these cells remained distinct from the conjunctival melanocytes (*KIT*+ and LEF1+) (Fig. 5M-P).

#### Macrophages

Although all macrophages grouped together in a single cluster regardless of tissue source, closer inspection indicated substantial heterogeneity. Macrophages of corneal origin were too few in number to form a distinct cluster, but segregated within a discrete zone, demonstrating expression of *CXCR4* and *ITGA4* but not *LYVE1* or *F13A1*. In contrast, the remainder of the macrophage cluster, which derived from the other tissues, expressed *LYVE1, STAB1* and *F13A1* (Fig. 5G, *SI Appendix*, Fig. S3D).

### Disease Associations

Many genes have been implicated in susceptibility to ocular diseases that result from defects in the anterior segment. We explored expression patterns of 924 disease-associated genes to the cell types in our atlas. Simultaneous interrogation of all anterior segment cell types provides insight into pathophysiology because the broad range and variability of phenotypes observed in many diseases may reflect variable contribution of multiple tissues. Moreover, even when the most striking phenotypes are observed in single ocular tissues, pathophysiological changes in surrounding tissues may contribute to disease state.

### Mendelian Genes

#### Anterior Segment Dysgenesis (ASD)

Congenital and developmental abnormalities of the anterior segment result from a complex interplay of developmental, embryological and genetic factors that lead to abnormalities of the iris, iridocorneal angle drainage structures, cornea, or combinations of these abnormalities (Ma 2019). Due to dysregulated aqueous outflow, approximately 50% of patients with ASD develop glaucoma (Alward 2000). Clinical subtypes of ASD include Axenfeld-Rieger Anomaly, Peters Anomaly, primary congenital glaucoma, and aniridia/iris hypoplasia. Genes implicated in ASD and glaucoma include *PAX6, PITX2, FOXC1, CYP1B1, LTBP2, FOXE3, PITX3, B3GLCT, COL4A1, PXDN, and CPAMD8*. Many of these genes, which play well-defined roles in ocular development, continue to be expressed in relevant adult cell types within the anterior segment (Fig. 6A,B *SI Appendix*, Fig. S5A).

**Figure 6.**
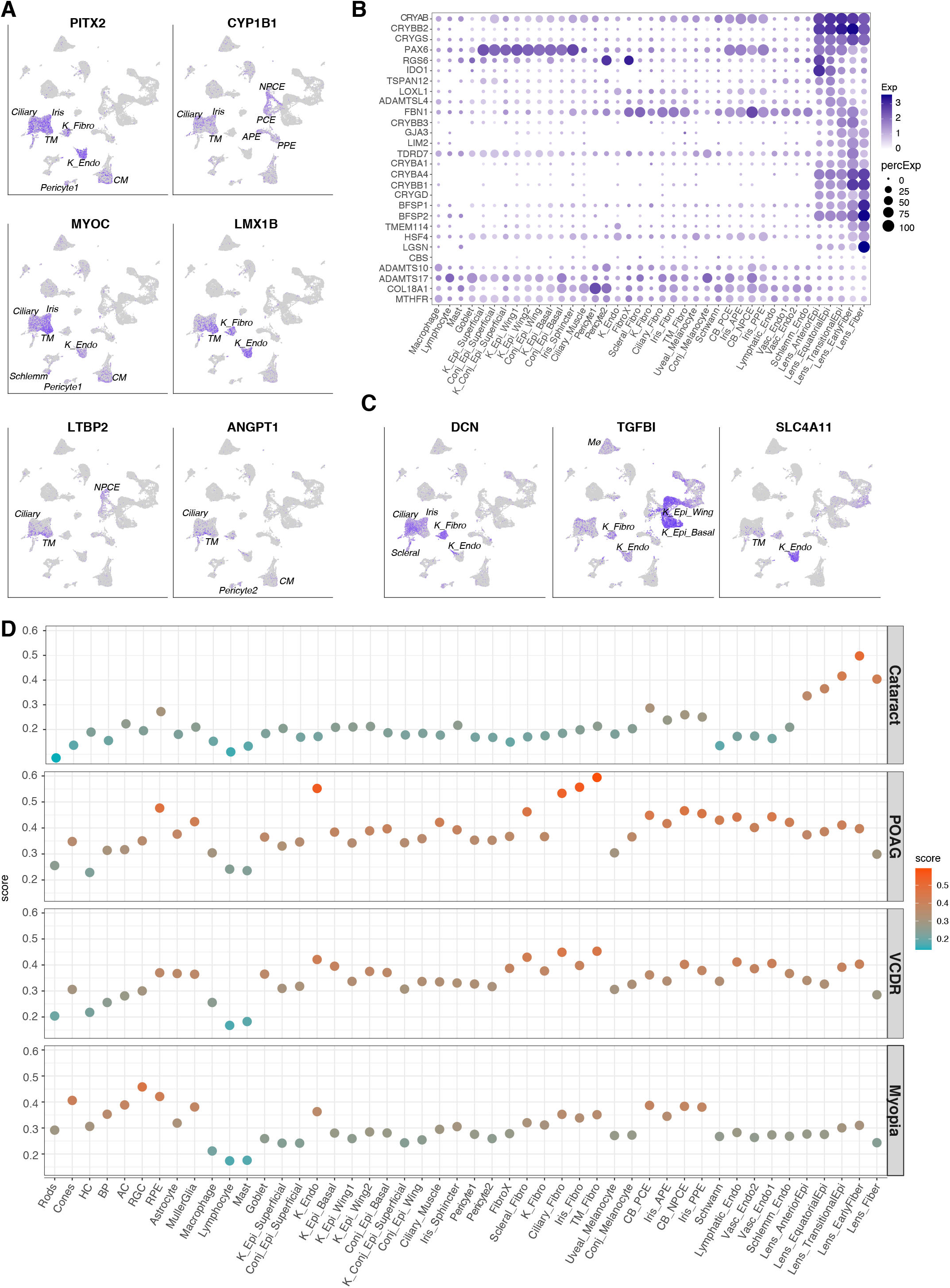
Expression of ocular disease-associated genes. **A** Feature plots demonstrating enriched expression of glaucoma-associated genes within cell types localized to the iridocorneal angle drainage structures and in some cases the ciliary epithelium. **B** Dot plot showing cataract- and ectopia lentis-associated genes. **C** Feature plot showing expression patterns of genes implicated in corneal dystrophies. **D** Dot plot showing cell-type specific enrichment scores of genes identified through GWAS for common ocular conditions or traits. Major retinal cell types from a normal macular sample are also included. Cell type abbreviations as in previous figures. POAG, primary open angle glaucoma; VCDR, vertical cup to disc ratio

#### Glaucoma

Glaucoma is a phenotypically heterogeneous disease with a final common outcome of optic nerve degeneration, with elevated IOP representing the only known modifiable risk factor (Stein et al JAMA 2021). We explored expression patterns of Mendelian genes associated with glaucoma (including ASD-related glaucoma) and elevated IOP, including *MYOC, ANGPT1, ANGPT2, LMX1B* as well as *PITX2, FOXC1, LTBP2*, and *CPAMD8*. As expected, most of these were robustly expressed in outflow pathway cells and other glaucoma-relevant cell types (Fig. 6A).

#### Cataract

Defined as opacity in the crystalline lens, cataract is the leading cause of blindness worldwide and generally follows a bimodal age distribution corresponding to congenital and age-related onset. Hereditary congenital cataracts, presenting at birth or during infancy, have been associated with highly penetrant genetic mutations in lens crystallins, growth factors, transcription factors, connexins, intermediate filament proteins, membrane proteins, the protein degradation apparatus, and a variety of other pathways including lipid metabolism (Shiels 2017). Crystallin genes were strongly expressed across lens epithelial and fiber cell types with gradient differences for a subset of genes. For example, *CRYAB, CRYGS* and *CRYBB2* were broadly expressed across all lens clusters (spanning both epithelial and fiber cell types), whereas *CRYBB3* and *CRYBA1* were most prominently expressed in transitional epithelium and early fiber cells, and *CRYBA4, CRYGD* and *CRYBB1* were enriched in lens fibers (Fig. 6B). The connexin gene, *GJA3*, demonstrated a similar expression pattern beginning in transitional epithelium and extending variably into the fiber cells. Among other associated congenital cataract genes, we noted peak expression either in the early fiber cells (e.g., *LIM2, TDRD7*) or the more mature fiber cells (e.g., *BFSP1, BFSP2, TMEM114, HSF4*). Only a minority of cataract-associated genes demonstrated predominant expression in the lens epithelial (as opposed to fiber cell) clusters, and most of these were in the setting of pleomorphic syndromes (*PAX6, RGS6, TSPAN12*).

#### Corneal Dystrophies

The corneal dystrophies are inherited conditions that impair the transparency of the cornea, leading to reduced vision and in some cases chronic ocular pain (Lisch 2020). While these conditions have variable presentations, many have clinical findings predominantly localized within one of the three major layers of the cornea. We found a tight correspondence between cell-type specific expression and the predominant clinical finding. Genes implicated in epithelial corneal dystrophies, including *KRT3* and *KRT12* (Meesman) and *COL17A1* (Recurrent Erosion), were expressed predominantly in the wing and basal cells of the corneal epithelium (*SI Appendix*, Fig. S2B). Genes implicated in endothelial dystrophies, including *COL8A2* (Posterior polymorphous), and *SLC4A11* (Congenital Hereditary type 2), were expressed predominantly in the corneal endothelium (Fig. 6C). Finally, genes implicated in stromal dystrophies, including *CHST6* (Macular), *DCN* (Congenital Stromal), and *TGFBI* (Lattice and Granular), were expressed in keratocytes. DCN and TGFBI were also expressed in other corneal types, consistent with their more varied phenotype (Fig. 6C).

#### Ectopia Lentis (EL)

is a condition in which the crystalline lens is abnormally positioned within the eye, most often due to damaged, dysfunctional or absent zonular fibers (Chandra et al 2014, Bassnett 2020). The most common cause of EL is trauma but mutations have been implicated in both isolated cases (e.g., *ADAMTSL4*) and in syndromes such as Marfan Syndrome (*FBN1*), Weill Marchesani Syndrome (*ADAMTS10, ADAMTS17*) Knobloch Syndrome (*COL18A1*), Aniridia (*PAX6*), and Homocystinuria (*CBS, MTR, MTHFR*). EL-associated genes were consistently expressed in the non-pigmented ciliary epithelial and/or the equatorial and transitional lens epithelial cells, which are located in areas adjacent to zonules, suggesting that they contribute to ongoing structural integrity of this important apparatus (Fig. 6B). *LOXL1*, implicated in pseudoexfoliation syndrome – a condition in which zonular fibers become brittle over time – was also expressed most prominently in the equatorial lens epithelium (Fig. 6B).

### Susceptibility Genes Nominated by GWAS Analysis

Susceptibility to common ocular conditions such as myopia, age-related cataract and primary open angle glaucoma (POAG) is conferred by multiple common genetic variants with small effect sizes, many of which have been identified by Genome Wide Association Studies (GWAS). In addition to interrogating each gene individually (*SI Appendix*, Fig. S6A-H), we developed a gene expression score (see Methods), to explore whether GWAS-identified genes associated with these complex traits were more likely to be expressed in specific subsets of cell types. Because both the anterior and posterior segments are involved in POAG and myopia, we applied this analysis to key retinal cell types, which we profiled recently (Yan et al., 2020), alongside the anterior segment atlas (Fig. 6D, *SI Appendix*, Fig. S5B). Patterns were broadly consistent with our current knowledge of disease mechanisms. Genes associated with cataract were enriched in lens cell types. Genes associated with POAG were most enriched in trabecular meshwork cells, cell types of outflow tissues within the iridocorneal angle (ciliary and iris fibroblasts), and corneal endothelium. Moderate enrichment scores in RPE, Muller glia, scleral fibroblasts, ciliary body epithelium and vessel endothelium are consistent with the disease’s complex pathophysiology and potential diversity of phenotypes. Genes associated with Vertical Cup-to-Disc Ratio (VCDR) demonstrated similar enrichment patterns to POAG, albeit less pronounced. Finally, myopia-associated genes are most strongly enriched in cells of the neural retina (specifically retinal ganglion cells, RPE and cones), as well as uveal epithelium and corneal endothelium.

## DISCUSSION

We used snRNAseq to profile cells comprising the human anterior segment, generating discrete cell atlases of the cornea, limbus, iris, ciliary body and lens. We then merged these datasets to perform an integrated analysis that enabled holistic appraisal of these cells across contiguous and interdependent ocular tissues. Finally, we used the atlases to interrogate cell-type specific expression patterns of disease-associated genes, gleaning insights into where they act.

### Tissue-specific and Shared Cell Types

Analysis of anterior segment structures defined many clusters, each presumably belonging to a single cell type or a small number of closely related types that were too transcriptomically similar to separate by unsupervised methods in the current dataset. They are presented in Figures 1 (cornea), 2 (limbus), 3 (iris and ciliary body) and 4 (lens), along with histological results (immunohistochemistry *and in situ* hybridization) to validate the assignment and determine the localization of many of them. Adding these results to those from our recent analysis of the trabecular meshwork and contiguous structures (van Zyl et al., 2020), but taking “shared” types from the integrated analysis into account, we estimate that there are at least 40 cell types in the human anterior segment. Our results are generally concordant with those of the recent “whole eye” atlas of Gautam et al. (2021), but the larger number of cells in our dataset than theirs (191,992 vs 50,000 total, of which only a minority were from anterior segment) allowed us to document far more cell types.

It was apparent that some of the types isolated from a particular tissue were very similar to ones isolated from other tissues. This is unsurprising for three reasons. First, although the structures we analyzed can be roughly separated by dissection, some are contiguous without sharply defined borders. For example, the iris root, trabecular meshwork and ciliary muscle share a common insertion area at the iridocorneal angle (Fig. 1B) and limbus is continuous with the cornea and sclera (Fig. 2A) such that discrete anatomical dissection at the cellular level is impossible. Thus, some contamination is inevitable, and caution must be taken when interpreting the “tissue source” of a given cell type. Second, some of the major cell types, such as macrophages, are motile and can easily cross tissue boundaries. Third, some types, such as melanocytes, Schwann cells and outflow fibroblasts populating the iridocorneal angle, are derived from the neural crest, a migratory population that invade multiple structures during development (Wang Rattner and Nathans, 2021).

By combining transcriptomes from contiguous tissues, reclustering them, and visualizing key markers with histological methods, we were able to determine which types are shared and which are primarily confined to specific tissues. Most epithelial types were tissue-specific whereas cells derived from the blood (mast cells, macrophages and lymphocytes) and Schwann cells (crest-derived) were shared. For other groups, distribution was more complex. For pericytes, melanocytes and endothelia, one type populated multiple tissues whereas the other (or others for endothelia) were tissue specific. Of particular interest were the melanocytes, with a conjunctival type derived from the corneoscleral wedge on the surface of the eye and a uveal type inside the eye present in multiple structures. The substantial differences between their transcriptomes may carry important implications in disease research. Uveal melanoma and conjunctival melanoma – representing serious oncologic diagnoses – behave very differently in the clinical setting and consequently require different treatment approaches (Harbour, 2016); an improved understanding of the molecular differences between them may provide insight into pathogenesis and therapeutic strategies.

Additional insights were gleaned for fibroblasts. Despite representing a diverse group of cells that play a key role in maintaining health and driving disease, fibroblasts throughout the body have historically been poorly characterized. Single-cell technologies have contributed significantly to advancing our understanding of their heterogeneity and family relationships as well as their contribution to disease states (Buechler et al., 2021). Our current work extends this analysis to fibroblasts within the anterior segment of the eye, particularly those populating the glaucoma-relevant tissue of the trabecular meshwork. Indeed, only by examining the trabecular meshwork tissue within its larger context of surrounding contiguous tissues were we able to appreciate both its relatedness to other neighboring tissues including the cornea, ciliary body and iris as well as its unique aspects. Among the anterior segment fibroblast cells, we found the TM cells to be most closely related to fibroblasts residing in the ciliary body and iris tissues, with certain unique markers as well as other more common markers with gradient expression patterns. In contrast, fibroblasts from the cornea and sclera were distinctly different, each defined by a unique set of ECM and collagen genes. In addition to the TM, both the iris stroma and ciliary muscle are exposed to aqueous humor, with the latter providing it with an alternative egress path. While similarities in these cells’ gene expression could relate to them potentially deriving from a common neural crest population (Czvekl & Tamm 2004), another factor may be related to their shared exposure to aqueous outflow.

A prominent illustration of tissue-specific cell types exhibiting relationships corresponding to their contiguity and embryological origins involved the cells populating the uveal epithelial bilayer lining the iris and ciliary body, whose origins trace to the neuroectodermal bilayer of the optic cup (Fuhrmann, 2010). The iris posterior pigmented epithelium demonstrated closest transcriptomic relation to the ciliary body non-pigmented epithelium, together comprising the inner layer of the uveal epithelial bilayer derived from the inner layer of the embryological optic cup. Similarly, the iris anterior pigmented epithelium demonstrated closest relation to the ciliary body pigmented epithelium, together comprising the outer layer of the uveal epithelial bilayer derived from the outer layer of the embryological optic cup. While we did not integrate posterior segment tissues into our analysis, we speculate that retinal pigment epithelium may show similarity to the other two cell types in this outer uveal epithelial layer. Similarly, while the neural retina ultimately takes on exquisite complexity, its embryological origin links it to the inner uveal layer, which may account for some residual neural-type expression patterns in the ciliary body non-pigmented epithelium and the iris posterior pigmented epithelium (e.g., *CRB1, LRRN2, CACNA1E*).

A caveat to the classification system is that we defined clusters or types using clustering algorithms that, while nominally unsupervised, are not entirely objective. This is especially true in cases where gene expression differences occur on a continuum among cells. For example, in several instances we noted that expression of specific genes displayed gradients within a cluster, suggesting the existence of either additional closely related cell types or cell state-related patterns of expression. Conversely, given the limited numbers of cells used, there may be tissue-specific differences among nominally “shared” types that we were unable to detect.

### Disease-associated Genes

Our cell atlas of the human anterior segment provides a valuable resource for analyzing genes implicated in ocular disease, particularly when combined with our human retinal cell atlas (Yan et al., 2020). First, it expands our insight into the complexity of glaucoma, where many genes previously shown to be expressed in TM (van Zyl et al, 2020) are also expressed in other relevant tissues including the iris, ciliary body and cornea. For example, outflow fibroblasts (including those populating the TM, ciliary body and iris), corneal endothelium and vessel endothelia (including those populating Schlemm canal, lymphatic and blood vessels) emerged as key cell types expressing genes associated with POAG and IOP. Second, it allows further exploration of other anterior segment diseases. For example, in cataract, transitional lens epithelium and early fiber cells were strongly overrepresented among genes associated with cataract; in myopia, the uveal epithelium, RPE and various neural cell types in the retina appeared to stand out as important expressors of associated genes.

## CONCLUSION

A cell atlas of the human eye’s anterior segment was generated by integrating the transcriptomes of 191,992 single nuclei derived from 6 key structures including the cornea, limbus, iris, ciliary body and lens. Beyond highlighting both the diversity and relatedness of cell types populating this complex sensory organ, this atlas facilitates direct interrogation of disease-relevant gene expression patterns across individual cell types. We hope this atlas will serve as a valuable resource to fuel current and future investigations in ophthalmology and vision science.

## MATERIALS & METHODS

### Tissue Acquisition, Dissection and Processing

Human ocular tissues were obtained post-mortem at a median of 6 hours from death either from Massachusetts General Hospital (MGH) via the Rapid Autopsy Program or from The Lion’s Eye Bank in Murray, Utah. Whole globes were transported to either to Harvard University or the University of Utah in a humid chamber on ice and processed within an hour of enucleation.

Dissection was performed under a surgical microscope. To isolate the anterior segment, a surgical blade was used to make a small stab incision ∼4 mm posterior to the limbus at the pars plana. The incision was extended circumferentially with curved scissors to yield a separated anterior segment and posterior segment. Each segment was placed in a petri dish filled with AMES’ medium equilibrated with 95% O_2_/5% CO_2,_ or sterile PBS, on ice.

The anterior segment was further dissected into its component tissues. The lens was liberated from the ciliary body by lysing the zonular attachments with curved microscissors. The lens capsule and superficial lens cells were isolated by making a small curvilinear incision in the anterior lens capsule with a surgical blade, and prolapsing the lens nucleus and cortical material, which was then discarded. The ciliary body was separated from the iris via blunt dissection with forceps and was then divided into 4 equal parts. The iris was divided similarly. After trephination of the central cornea using a 6 mm corneal punch, the remaining corneoscleral rim was addressed. Any large portions of ciliary muscle left behind after blunt dissection of the iris and ciliary body were trimmed off with microscissors. The trabecular meshwork was gently freed from the corneoscleral rim, often in multiple strips, by inserting one prong of the microforceps into Schlemm’s canal and gently clamping onto the trabecular meshwork tissue with the other prong and applying steady traction. The remaining corneoscleral tissue was cut into wedges and each wedge was trimmed to include approximately 1-2 mm of peripheral cornea and 1-2 mm of perilimbal sclera.

For snRNA-seq, each dissected tissue was immediately placed along the wall of a cryogenic vial, ensuring minimal liquid was present, and submerged in dry ice or liquid nitrogen. For long-term storage, the cryogenic vials were moved to a -80°C freezer. Samples collected at the University of Utah were transferred to Harvard University via overnight shipping on dry ice. For single-nuclei isolation, frozen tissue samples were homogenized in a dounce homogenizer with NP-40 (0.1%) Tris-based lysis buffer and passed through a 40-µm cell strainer. The filtered nuclei were then pelleted at 500 rcf over 5 min and resuspended in 2% BSA with DAPI counterstain for cell sorting on a flow cytometer. The sorted nuclei were pelleted again at 500 rcf over 5 min and resuspended in 0.04% non-acetylated BSA/PBS solution and adjusted to a concentration of 1000 nuclei/µL. The integrity of the nuclear membrane and presence of non-nuclear material were assessed under a brightfield microscope before loading into a 10X Chromium Single Cell Chip with a targeted recovery of 6000 nuclei.

Single nuclei libraries were generated with either Chromium 3’ V3, or V3.1 platform (10X Genomics, Pleasanton, CA) following the manufacturer’s protocol. Briefly, single nuclei were partitioned into Gel-beads-in-EMulsion (GEMs) where nuclear lysis and barcoded reverse transcription of RNA would take place to yield full-length cDNA; this was followed by amplification, enzymatic fragmentation and 5’ adaptor and sample index attachment to yield the final libraries.

### RNA Sequencing and Data Analysis

Libraries were sequenced on Illumina HiSeq 2500 at the Broad Institute or NovaSeq at the Bauer Core Facility at Harvard University. The single nuclei RNA sequencing data were demultiplexed and aligned using Cell Ranger software (version 4.0.0, 10X Genomics, Pleasanton, CA). Reads were aligned to the human genome GRCh38 version 101 with the following modifications: 1. Human genome files GRCh38 were downloaded from Ensembl; 2. the “transcript” lines in the .gtf file were converted into “exon” lines without changings to the remaining; 3. The reference files were generated using the cellranger “mkref” function; 4. Reads were aligned to the modified file and counted using cellranger “count” function with the chemistry option specified as “SC3Pv3”.

The downstream analysis pipeline was similar to that in van Zyl et al. with the exception that the R package “Seurat” (version 4.0.3) was used. Data from each tissue region was first analyzed separately. Briefly, the count matrix was filtered so only cells with more than 1000 genes detected were included. The dataset was then split into batches based on the donors of the samples and normalized individually using SCTransform. The top 2000 features common across batches were used to identify anchors for data integration. It was then scaled, and principal component (PC) analysis was performed. The top 50 PCs were selected for visualization and clustering. An array of different resolutions was assessed when clustering. Starting with the high resolutions, clusters were evaluated based on number of differentially expressed (DE) genes between closely related clusters. The clustering with maximal resolutions and strong DE genes (more than five DE genes on both up- or down-regulated side between closely related clusters) were, in most cases, taken as the putative types. However, in some cases as pointed out in the text, sub-clustering among certain population was identified only with the higher resolution, and that was projected into the final clustering result. DE test was performed using the “MAST” method. Transcriptomic relationship among clusters was built using the scaled data after integration.

For the anterior segment integrated analysis, after clustering of cells, a downsampling procedure was introduced so a maximum of 1000 cells from each putative type in each tissue were merged into a combined dataset and re-analyzed. Only contiguous tissues that were spatially and transcriptomically close to each other were included; this excluded the lens. In assessing the expression of disease related genes in the eye, both the lens and from retinal datasets were subsequently included for completeness. The disease gene lists were generated from the following sources: Disc Diameter and VCDR from Han et al 2021; POAG from Gharakhani et al 2021; IOP from Khawaja et al 2018; Myopia from Hysi et al 2020; Astigmatism from the National Human Genome Research Institute-European Bioinformatics Institute (NHGRI-EBI) GWAS catalog (Buniello et al 2019, downloaded 20 February 2021); and Cataract from the Cat-Map database filtered for genes with documented human phenotypes (Shiels et al 2010; https://cat-map.wustl.edu/). Only genes that were expressed in more than 25% of cells in any cluster were included in the analysis. To visualize the expression of each disease gene set, a gene set score was calculated as follows: 1. The genes in the gene set list were first filtered so only those that expressed in more than 25% of cells in any putative type were assessed. 2. For each cell i, the mean expression value of the genes in the set j (Exp_(i,j)) and that of the total transcripts (^−^(Exp_i)) were calculated. 3. The score of gene set j in cell i (S_(i,j)) was calculated as S_(i,j)=Exp_(i,j)-^−^(Exp_i). 4. The averaged score of gene set j in each putative type was visualized in Figure 6D and *SI Appendix* Fig. S5B.

### Histology

Corneoscleral wedges or whole globes were fixed in 4% paraformaldehyde in PBS for 2-24 hr and then transferred to PBS. The fixed samples were sunk in 30% sucrose in PBS overnight at 4°C, then embedded in tissue freezing medium and mounted onto coated slides in 10-50 μm meridional sections with ProLong Gold Antifade Mountant. For immunohistochemistry, slides were incubated for 1 hour in 5% donkey serum and 0.3% TritonX block at room temperature, overnight with primary antibodies in Renoir Red Diluent at 4°C, and 2 hours with secondary antibodies in 3% donkey serum and 0.3% TritonX at room temperature. Antibodies used are listed in **Supplementary Table 3**. Single molecule fluorescent *in situ* hybridization was performed using commercially available RNAScope Multiplex Fluorescent Assay V2 (Advanced Cell Diagnostics, Newark, CA). Briefly, slides were baked in a HybEZ oven (Advanced Cell Diagnosics, Newark, CA) at 60°C for 30 minutes, then tissue freezing medium was removed by soaking the slides in sterile PBS. A 1-hour post-fix in 4% PFA was performed at 4°C and the slides were then dehydrated in serial ethanol concentration rinses. The slides were incubated for 10 minutes at room temperature with the hydrogen peroxide provided in the assay kit diluted 1:1 in water then for 30 minutes with protease III at 40°C. Probe hybridization and subsequent steps were per standard manufacturer protocol. Probes used are listed in **Supplementary Table 3**.

### Image acquisition, processing and analysis

Images were acquired on Zeiss LSM 710 confocal microscopes with 405, 488-515, 568, and 647 nm lasers, processed using Zeiss ZEN software suites, and analyzed using ImageJ (NIH). Images were acquired with 16X, 40X or 63X oil lenses at the resolution of 1024×1024 pixels, a step size of 0.5-1.5μm, and 90μm pinhole size.

## Supporting information

van Zyl et al., Supplementary Figures and Tables

## ACKNOWLEDGEMENTS

This work was supported by NIH (5K12EY016335, EY028633 and U01 MH105960), the Chan-Zuckerberg Initiative (CZF-2019-002459) and the Klarman Cell Observatory of the Broad Institute of MIT and Harvard, an unrestricted grant from Research to Prevent Blindness to the Department of Ophthalmology and Visual Sciences, University of Utah, and by charitable donations to the Sharon Eccles Steele Center for Translational Medicine. We thank Chris Pappas and Lisa Nichols for their efforts to acquire and process ocular tissues, and the tissue donors and their families for their generosity.

