## Supplementary material for "Cell atlas of the human ocular anterior segment: Tissue-specific and shared cell types": van Zyl et al., Supplementary Figures and Tables

5 Equal contribution

\

**SI Appendix**

**Fig. S1. Distribution of cell types within each tissue**
Histograms show percentage of cells assigned to each type in each tissue
**A** Central cornea
**B** Internal and external limbus
**C** Limbal ocular surface epithelium (subset of B)
**D** Iris
**E** Lens
**F** Ciliary body

**Fig. S2. Cell types of the Internal and External Limbus**
**A** Clustering of 10,510 single-nucleus transcriptomes derived from corneoscleral wedge tissue that were identified as ocular surface epithelial cells. Data visualized by Uniform Manifold Approximation and Projection (UMAP).
**B** Violin plot showing common and selectively expressed genes in corneal and conjunctival epithelial cells.
**C** Composite image of limbal region with dashed lines approximating regions of the cornea, limbus and conjunctiva with underlying sclera. Boxed area is magnified to show LGR6-positive (red) conjunctival basal epithelial cells and MECOM-positive wing and superficial cells visualized with fluorescent RNA *in situ* hybridization.
**D** Dot plot showing expression of LGR6 and MECOM within cells of the limbus.
**E** Feature plots showing expression of genes associated with mitotically active and limbal progenitor cells. Pink dashed circle identifies putative transit-amplifying cells as an island within the corneal basal epithelium cluster; blue dashed circle identifies putative limbal progenitor cells within the conjunctival basal epithelium cluster.
**F** Clustering of 51,306 single-nucleus expression profiles derived from corneoscleral wedge (CSW) tissue visualized by UMAP. This panel shows contributions of cell types from the TM and CSW, which were processed separately and pooled for subsequent analysis.
**G** Feature plots showing expression of a selection of genes discussed in a previous analysis of TM (van Zyl et al., 2020).
**H** Fluorescent RNA *in situ* hybridization for *PPP1R1B* (red) and *TMEFF2* (green) highlights expression within cells of the TM as well as the ciliary muscle.
**I** Fluorescent RNA *in situ* hybridization for *C7* (red) and *BMP5* (green) highlights expression within cells at the base of the TM and most prominently within the ciliary muscle.
**J** Fluorescent RNA *in situ* hybridization for *NEB* (red) highlights cells confined to the TM and *PI16* (green) highlights those within the ciliary body near the scleral interface.
**K** Genes associated with Aqueous Veins / Collector channels, *SELE*, *ACKR1*, *COL15A1* (van Zyl et al 2020) co-cluster in the upper lobe of the Vasc_Endo cluster.
Mø, macrophage; Conj, conjunctival; Vasc, vascular; Endo, endothelium, Fibro, fibroblast; K_Endo, corneal endothelium; K_Scleral_Fibro, corneoscleral fibroblast; TM, Trabecular Meshwork; Conj_Epi, conjunctival epithelium; K_Epi, corneal epithelium; DAPI, 4′,6-diamidino-2-phenylindole.
Scale bars show 100μm

**Fig. S3. Integrated Analysis – Tissue and Donor Contributions**
**A** Contributions of donor cells to each cluster as visualized by Uniform Manifold Approximation and Projection (UMAP) with individual colors representing unique donor source. Due to superposition effects, not all dots are visible.
**B** Contributions of cells to integrated analysis by tissue source as visualized by Uniform Manifold Approximation and Projection (UMAP).
**C** Tissue sources of melanocyte cell types. All tissues except cornea contribute MET+ uveal melanocytes whereas KIT+ conjunctival melanocytes are exclusively derived from the CSW tissue.
**D** Tissue sources of macrophages. Macrophage-type cells in the cornea and corneal portion of CWS are LYVE-negative.

**Fig. S4. Integrated Analysis – Gene Expression and Histology**
**A** Dot plot showing differentially expressed marker genes for each cell type in the integrated analysis.
**B** Feature plots showing expression patterns of unique and shared genes within the fibroblast clusters. See Figure 5A for names of fibroblast types
**C** Fluorescent RNA *in situ* hybridization for *NELL2* (red) and *UNC5D* (green) highlights expression within TM fibroblasts confined to the trabecular meshwork
**D** Fluorescent RNA *in situ* hybridization for *TMEFF2* (red) highlights cells along the uveal border of the TM extending into the ciliary muscle and *ANGPTL7* (green) highlights those within the corneoscleral stroma and TM enriched in the juxtacanalicular region.
**E** Fluorescent RNA *in situ* hybridization for *TMEM178A* (green) and *NEB* (red) highlights cells within the TM.
**F** Immunostaining for PDPN and WIF1 shows iris fibroblasts most densely packed in the anterior border layer of the iris.
**G** Lymphatic endothelium within the conjunctival stromal expressing *PKHD1L1* (red), and vascular endothelium type 2 beneath the lymphatics expressing *BTNL9* (green); both visualized with RNA *in situ* hybridization.
**H** Fluorescent RNA *in situ* hybridization for *NDUFA4L2* (red) and *NOTCH3* (green) highlights expression of pericytes associated with vessels in the iris.
**I** Same image as in H, showing only the red channel representing *NFUFA4L2* expression.
**J** Fluorescent RNA *in situ* hybridization for *NDUFA4L2* (red) highlights pericytes associated with vessels in the conjunctival stroma.
**K** Same image as in J, with additional channel visualizing Fluorescent RNA *in situ* hybridization for *NOTCH3* (green).
Scale bars show 100μm

**Fig. S5. Human Disease Genes**
**A** Dot plot showing expression patterns of select genes implicated in anterior segment dysgenesis, cataract, glaucoma and ectopia lentis within the anterior segment and lens.
**B** Dot plot showing cell-type specific enrichment scores of genes identified through GWAS for common ocular conditions or traits. Major retinal cell types from a normal

macular sample are also included.  Similar plots for other disease groups are shown in Figure 6D.

**Fig. S6. Human Disease Genes**
Heatmap showing relative expression of genes implicated in susceptibility to ocular disease or in phenotypes related to disease. In most cases, genes were identified through GWAS, but some Mendelian genes are also included. The genes in the heatmap were used the expression scores for corresponding conditions or traits shown in Fig. 6D and S4B.
**A** Astigmatism
**B** Variations in optic disc diameter
**C** Elevated intraocular pressure
**D** Myopia
**E** Primary open angle glaucoma
**F** Refractive error
**G** Vertical cup to disc ratio
**H** Cataract

FIGURE S1

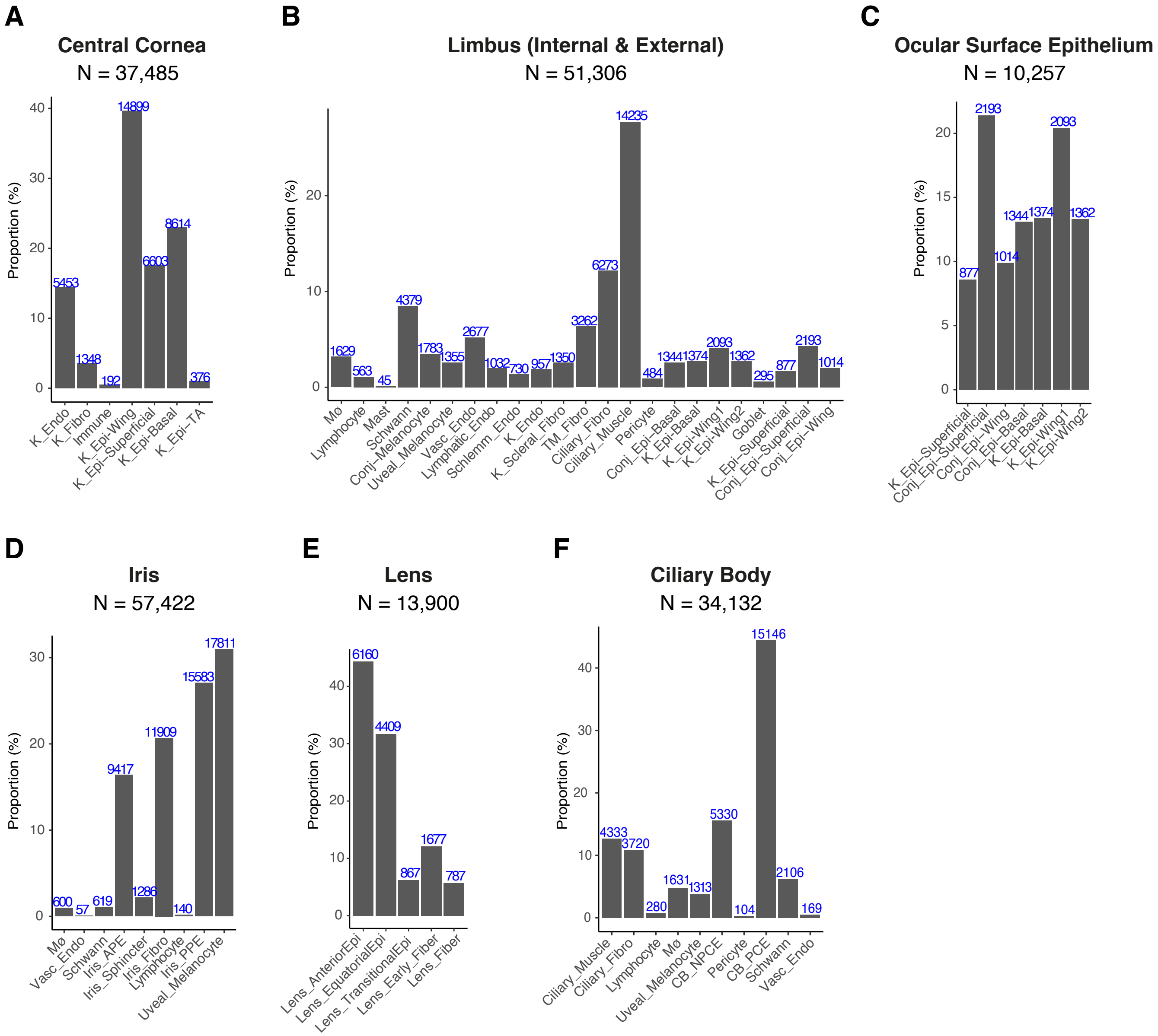

A
Central Cornea
N = 37,485
Proportion (%)
0
10
20
30
40
5453
1348
192
14899
6603
8614
376
K_Endo
K_Fibro
Immune
K_Epi-Wing
K_Epi-Superficial
K_Epi-Basal
K_Epi-TA
B
Limbus (Internal & External)
N = 51,306
Proportion (%)
0
10
20
1629
563
45
4379
1783
1355
2677
1032
730
957
1350
3262
6273
14235
484
1344
1374
2093
1362
295
877
2193
1014
Mø
Lymphocyte
Mast
Schwann
Conj_Melanocyte
Uveal_Melanocyte
Vasc_Endo
Lymphatic_Endo
Schlemm_Endo
K_Endo
K_Scleral_Fibro
TM_Fibro
Ciliary_Fibro
Ciliary_Muscle
Pericyte
Conj_Epi-Basal
K_Epi-Basal
K_Epi-Wing1
K_Epi-Wing2
Goblet
K_Epi-Superficial
Conj_Epi-Superficial
Conj_Epi-Wing
C
Ocular Surface Epithelium
N = 10,257
Proportion (%)
0
5
10
15
20
877
2193
1014
1344
1374
2093
1362
K_Epi-Superficial
Conj_Epi-Superficial
Conj_Epi-Wing
Conj_Epi-Basal
K_Epi-Basal
K_Epi-Wing1
K_Epi-Wing2
D
Iris
N = 57,422
Proportion (%)
0
10
20
30
600
57
619
9417
1286
11909
140
15583
17811
Mø
Vasc_Endo
Schwann
Iris_APE
Iris_Sphincter
Iris_Fibro
Lymphocyte
Iris_PPE
Uveal_Melanocyte
E
Lens
N = 13,900
Proportion (%)
0
10
20
30
40
6160
4409
867
1677
787
Lens_AnteriorEpi
Lens_EquatorialEpi
Lens_TransitionalEpi
Lens_Early_Fiber
Lens_Fiber
F
Ciliary Body
N = 34,132
Proportion (%)
0
10
20
30
40
4333
3720
280
1631
1313
5330
104
15146
2106
169
Ciliary_Muscle
Ciliary_Fibro
Lymphocyte
Mø
Uveal_Melanocyte
CB_NPCE
Pericyte
CB_PCE
Schwann
Vasc_Endo

FIGURE S2

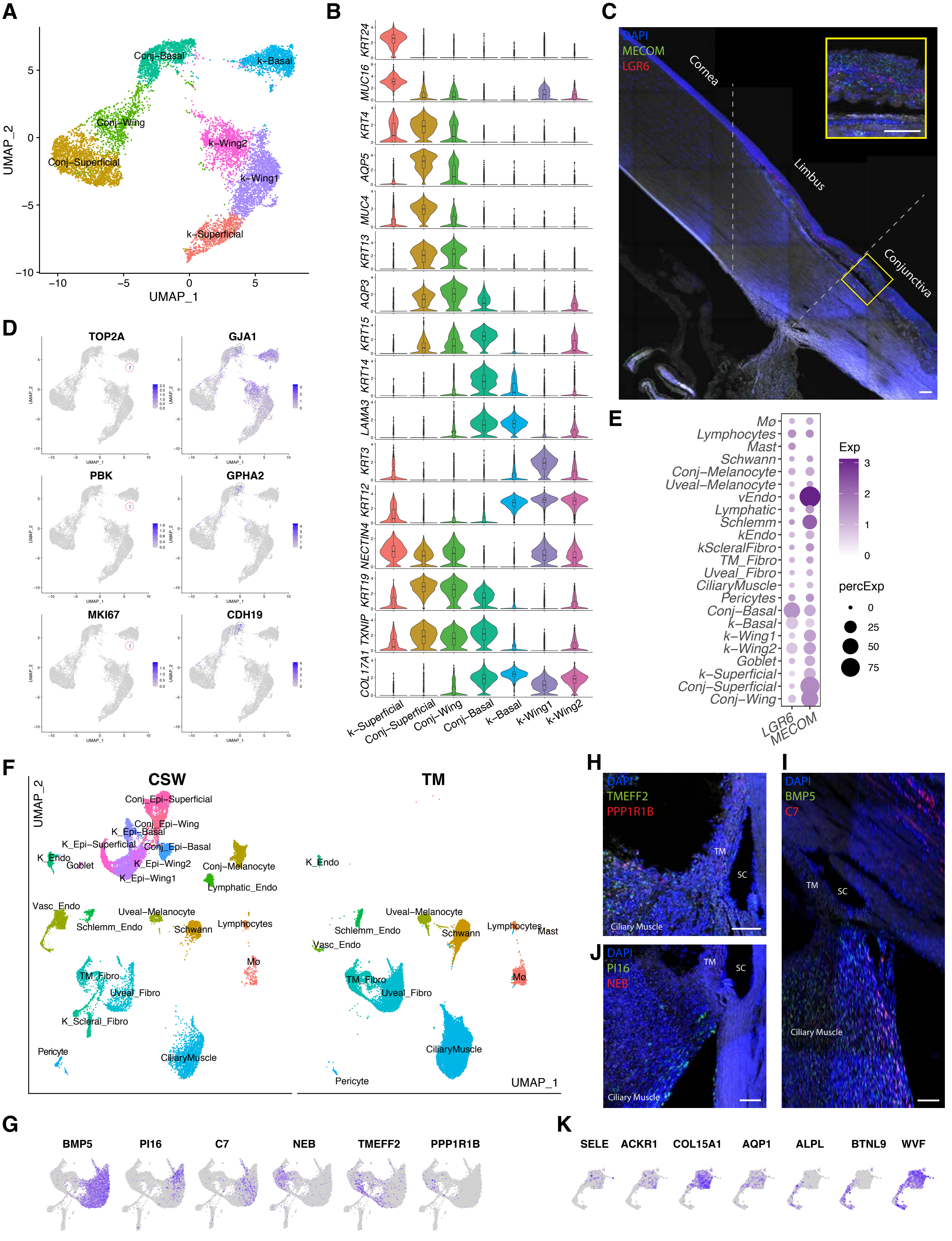

A
B
C
D
E
F
G
H
I
J
K
Conj–Basal
k–Basal
Conj–Wing
k–Wing2
k–Wing1
Conj–Superficial
k–Superficial
UMAP_1
UMAP_2
KRT24
MUC16
KRT4
AQP5
MUC4
KRT13
AQP3
KRT15
KRT14
LAMA3
KRT3
KRT12
NECTIN4
KRT19
TXNIP
COL17A1
DAPI
MECOM
LGR6
Cornea
Limbus
Conjunctiva
TOP2A
GJA1
PBK
GPHA2
MKI67
CDH19
Exp
percExp
CSW
TM
TMEFF2
PPP1R1B
BMP5
C7
PI16
NEB
SC
Ciliary Muscle
SELE
ACKR1
COL15A1
AQP1
ALPL
BTNL9
WVF

#### Figure S3

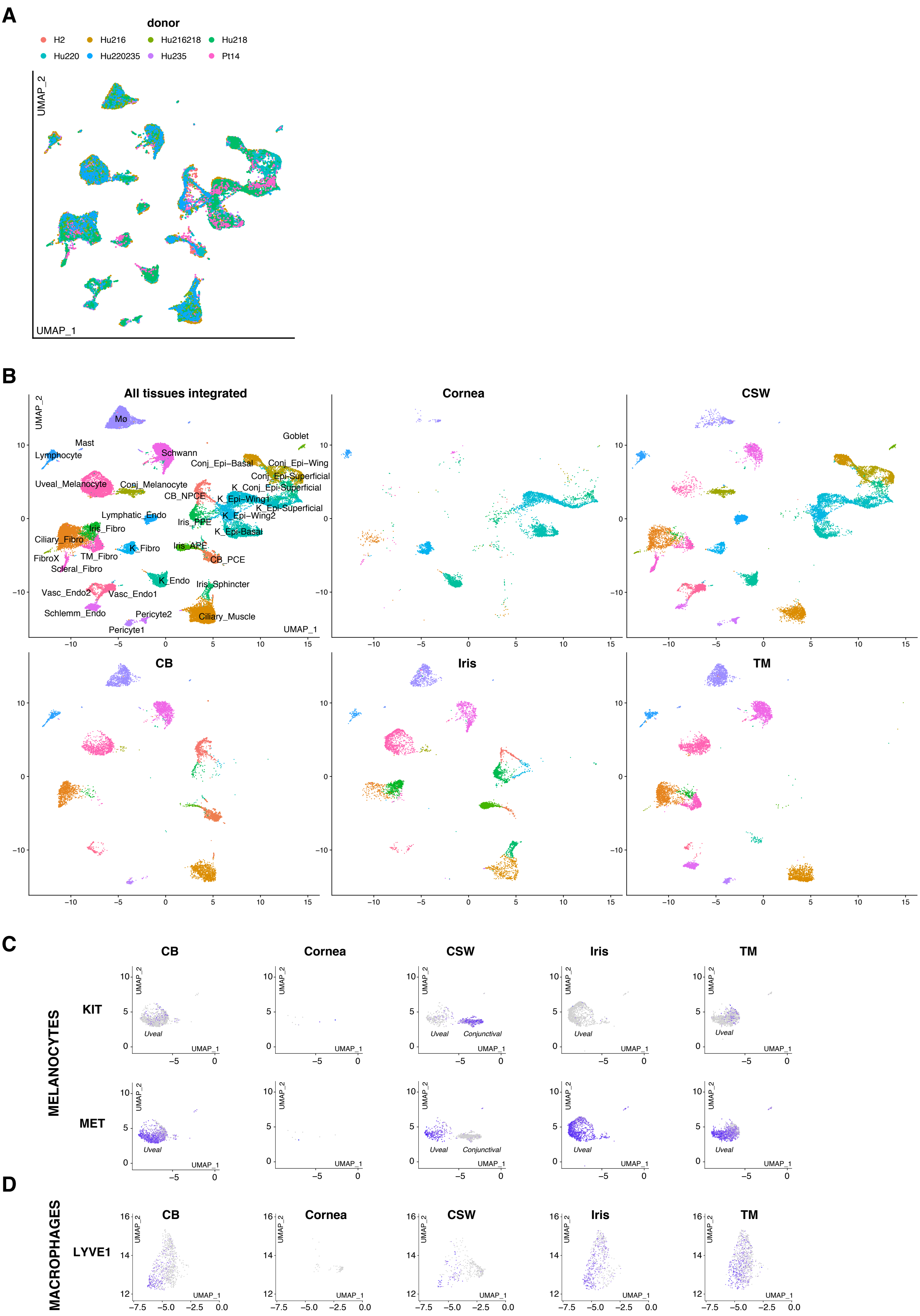
A
donor
H2
Hu216
Hu216218
Hu218
Hu220
Hu220235
Hu235
Pt14
UMAP_2
UMAP_1
B
All tissues integrated
Cornea
CSW
CB
Iris
TM
Mø
Mast
Lymphocyte
Schwann
Goblet
Conj_Epi−Basal
Conj_Epi−Wing
Conj_Epi-Superficial
Uveal_Melanocyte
Conj_Melanocyte
CB_NPCE
K_Conj_Epi-Superficial
K_Epi−Wing1
K_Epi-Superficial
K_Epi−Wing2
Lymphatic_Endo
Iris_PPE
Iris_Fibro
K_Epi-Basal
Ciliary_Fibro
K_Fibro
Iris_APE
FibroX
TM_Fibro
CB_PCE
Scleral_Fibro
K_Endo
Iris_Sphincter
Vasc_Endo2
Vasc_Endo1
Schlemm_Endo
Pericyte2
Ciliary_Muscle
Pericyte1
C
MELANOCYTES
KIT
MET
Uveal
Conjunctival
D
MACROPHAGES
LYVE1

### FIGURE S4

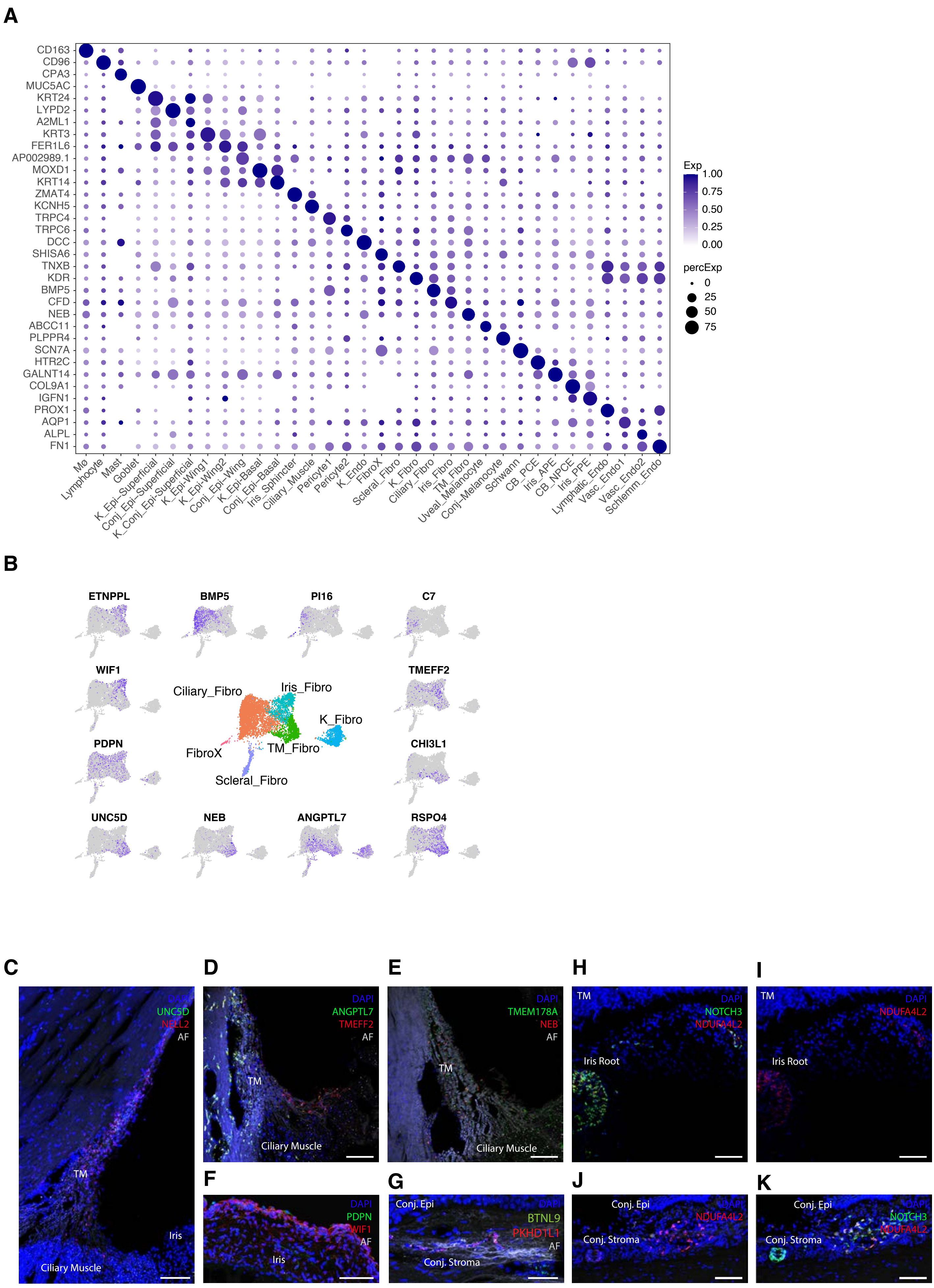
A
CD163
CD96
CPA3
MUC5AC
KRT24
LYPD2
A2ML1
KRT3
FER1L6
AP002989.1
MOXD1
KRT14
ZMAT4
KCNH5
TRPC4
TRPC6
DCC
SHISA6
TNXB
KDR
BMP5
CFD
NEB
ABCC11
PLPPR4
SCN7A
HTR2C
GALNT14
COL9A1
IGFN1
PROX1
AQP1
ALPL
FN1
Mo
Lymphocyte
Mast
Goblet
K_Epi-Superficial
Conj_Epi-Superficial
K_Conj_Epi-Superficial
K_Epi-Wing1
K_Epi-Wing2
Conj_Epi-Wing
K_Epi-Basal
Conj_Epi-Basal
Iris_Sphincter
Ciliary_Muscle
Pericyte1
Pericyte2
K_Endo
FibroX
Scleral_Fibro
K_Fibro
Ciliary_Fibro
Iris_Fibro
TM_Fibro
Uveal_Melanocyte
Conj_Melanocyte
Schwann
CB_PCE
Iris_APE
CB_NPCE
Iris_PPE
Lymphatic_Endo
Vasc_Endo1
Vasc_Endo2
Schlemm_Endo
Exp
1.00
0.75
0.50
0.25
0.00
percExp
0
25
50
75
B
ETNPPL
BMP5
PI16
C7
WIF1
TMEFF2
Ciliary_Fibro
Iris_Fibro
K_Fibro
FibroX
TM_Fibro
Scleral_Fibro
PDPN
CHI3L1
UNC5D
NEB
ANGPTL7
RSPO4
C
DAPI
UNC5D
NELL2
AF
TM
Iris
Ciliary Muscle
D
DAPI
ANGPTL7
TMEFF2
AF
TM
Ciliary Muscle
E
DAPI
TMEM178A
NEB
AF
TM
Ciliary Muscle
H
TM
DAPI
NOTCH3
NDUFA4L2
Iris Root
I
TM
DAPI
NDUFA4L2
Iris Root
F
DAPI
PDPN
WIF1
AF
Iris
G
Conj. Epi
DAPI
BTNL9
PKHD1L1
AF
Conj. Stroma
J
Conj. Epi
DAPI
NDUFA4L2
Conj. Stroma
K
Conj. Epi
DAPI
NOTCH3
NDUFA4L2
Conj. Stroma

#### FIGURE S5

**A**

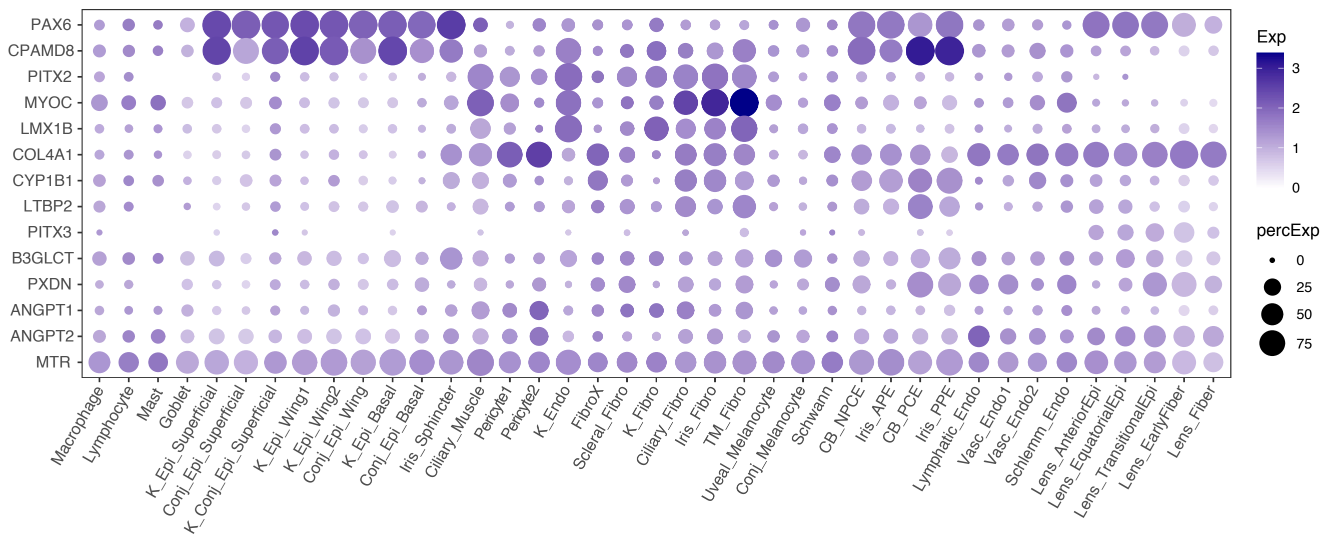

PAX6
CPAMD8
PITX2
MYOC
LMX1B
COL4A1
CYP1B1
LTBP2
PITX3
B3GLCT
PXDN
ANGPT1
ANGPT2
MTR
Macrophage
Lymphocyte
Mast
Goblet
K_Epi_Superficial
Conj_Epi_Superficial
K_Conj_Epi_Superficial
K_Epi_Wing1
K_Epi_Wing2
Conj_Epi_Wing
K_Epi_Basal
Conj_Epi_Basal
Iris_Sphincter
Ciliary_Muscle
Pericyte1
Pericyte2
K_Endo
FibroX
Scleral_Fibro
K_Fibro
Ciliary_Fibro
Iris_Fibro
TM_Fibro
Uveal_Melanocyte
Conj_Melanocyte
Schwann
CB_NPCE
Iris_APE
CB_PCE
Iris_PPE
Lymphatic_Endo
Vasc_Endo1
Vasc_Endo2
Schlemm_Endo
Lens_AnteriorEpi
Lens_EquatorialEpi
Lens_TransitionalEpi
Lens_EarlyFiber
Lens_Fiber
Exp
3
2
1
0
percExp
0
25
50
75


**B**

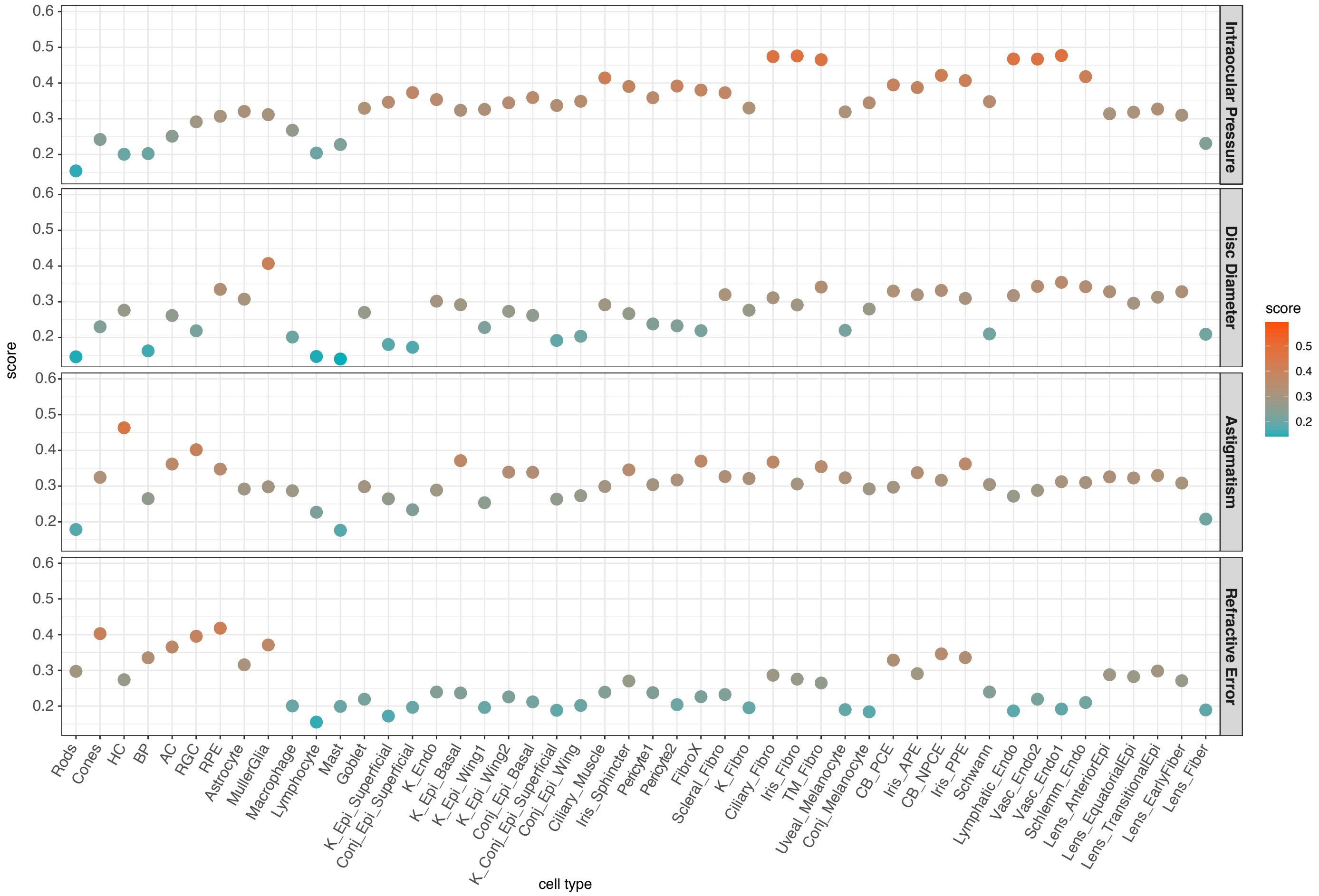

score
0.6
0.5
0.4
0.3
0.2
Intraocular Pressure
Disc Diameter
Astigmatism
Refractive Error
Rods
Cones
HC
BP
AC
RGC
RPE
Astrocyte
MullerGlia
Macrophage
Lymphocyte
Mast
Goblet
K_Epi_Superficial
Conj_Epi_Superficial
K_Endo
K_Epi_Basal
K_Epi_Wing1
K_Epi_Wing2
Conj_Epi_Basal
K_Conj_Epi_Superficial
Conj_Epi_Wing
Ciliary_Muscle
Iris_Sphincter
Pericyte1
Pericyte2
FibroX
Scleral_Fibro
K_Fibro
Ciliary_Fibro
Iris_Fibro
TM_Fibro
Uveal_Melanocyte
Conj_Melanocyte
CB_PCE
Iris_APE
CB_NPCE
Iris_PPE
Schwann
Lymphatic_Endo
Vasc_Endo2
Vasc_Endo1
Schlemm_Endo
Lens_AnteriorEpi
Lens_EquatorialEpi
Lens_TransitionalEpi
Lens_EarlyFiber
Lens_Fiber
cell type
score
0.5
0.4
0.3
0.2

### FIGURE S6A - Astigmatism

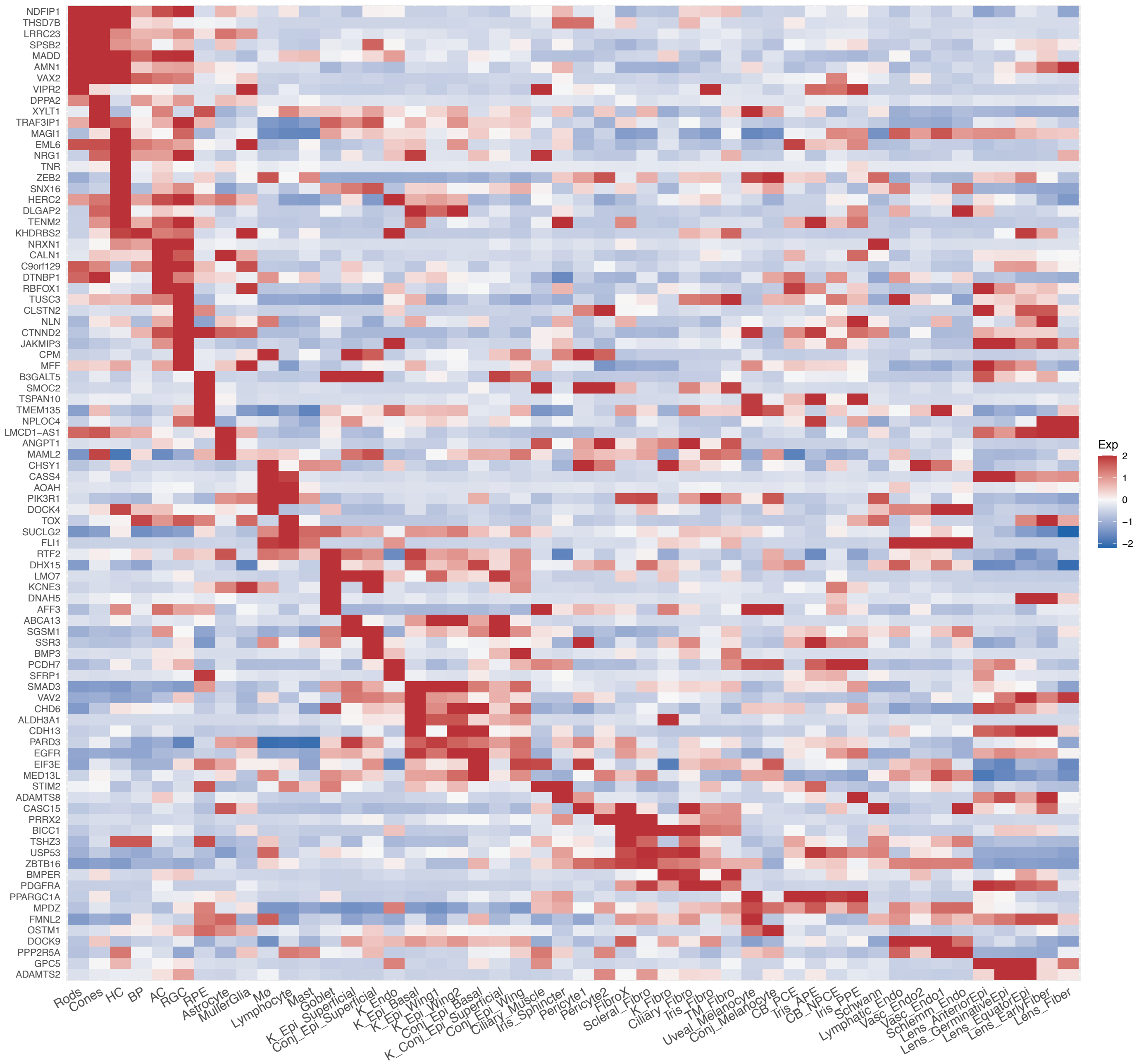

### FIGURE S6B - Disc Diameter

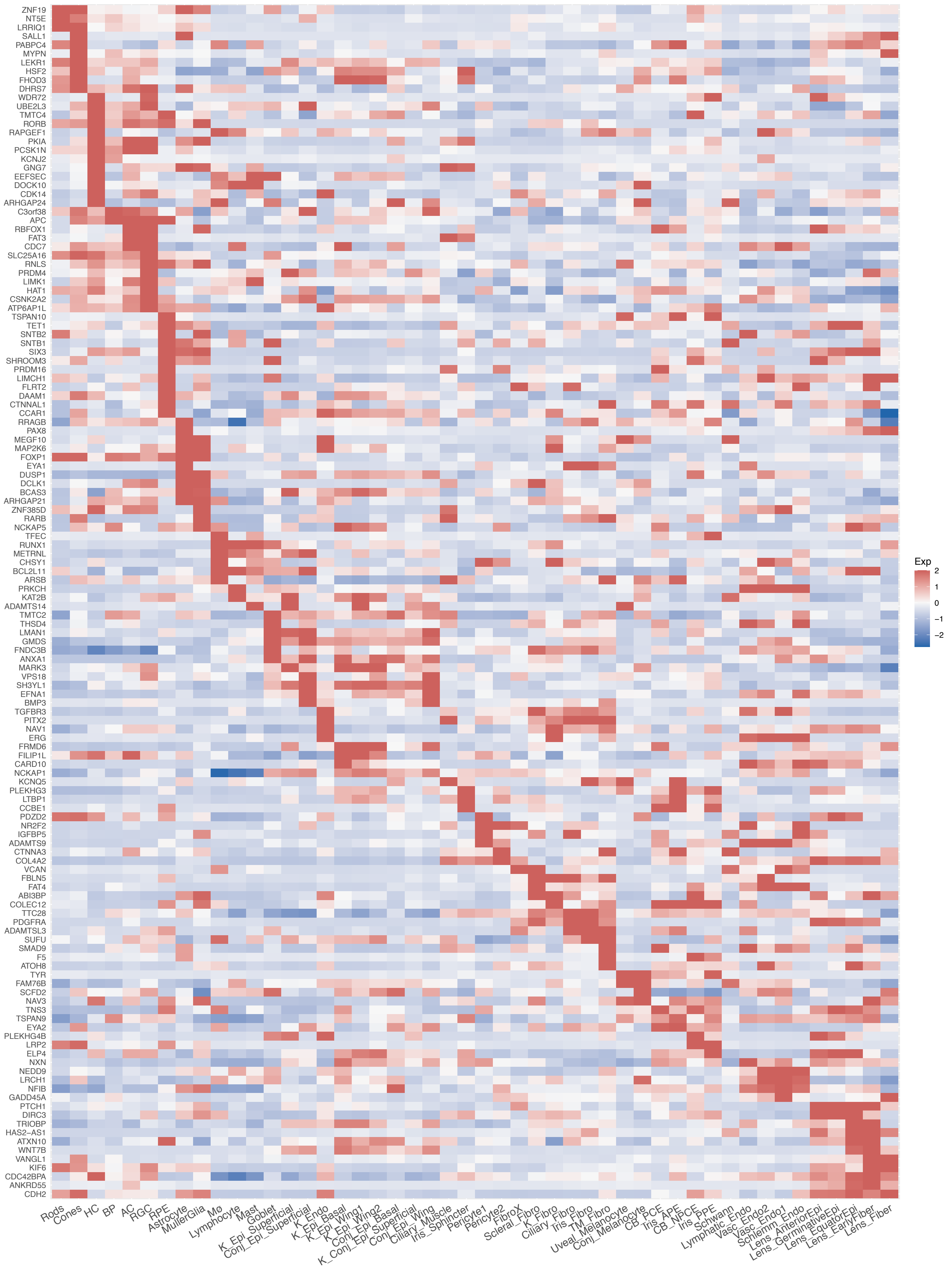

### FIGURE S6C - Intraocular Pressure (IOP)

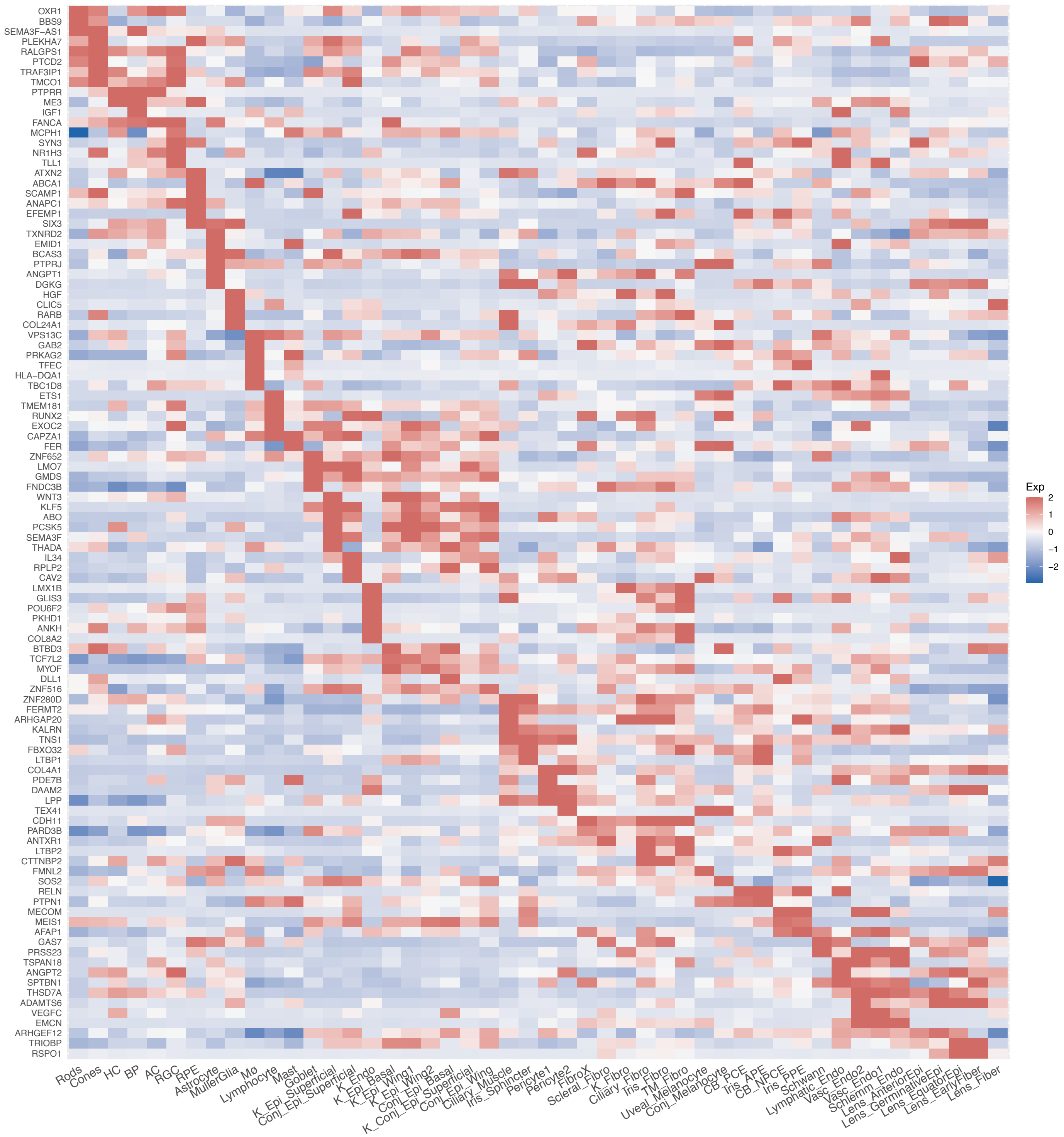

### FIGURE S6D - Myopia

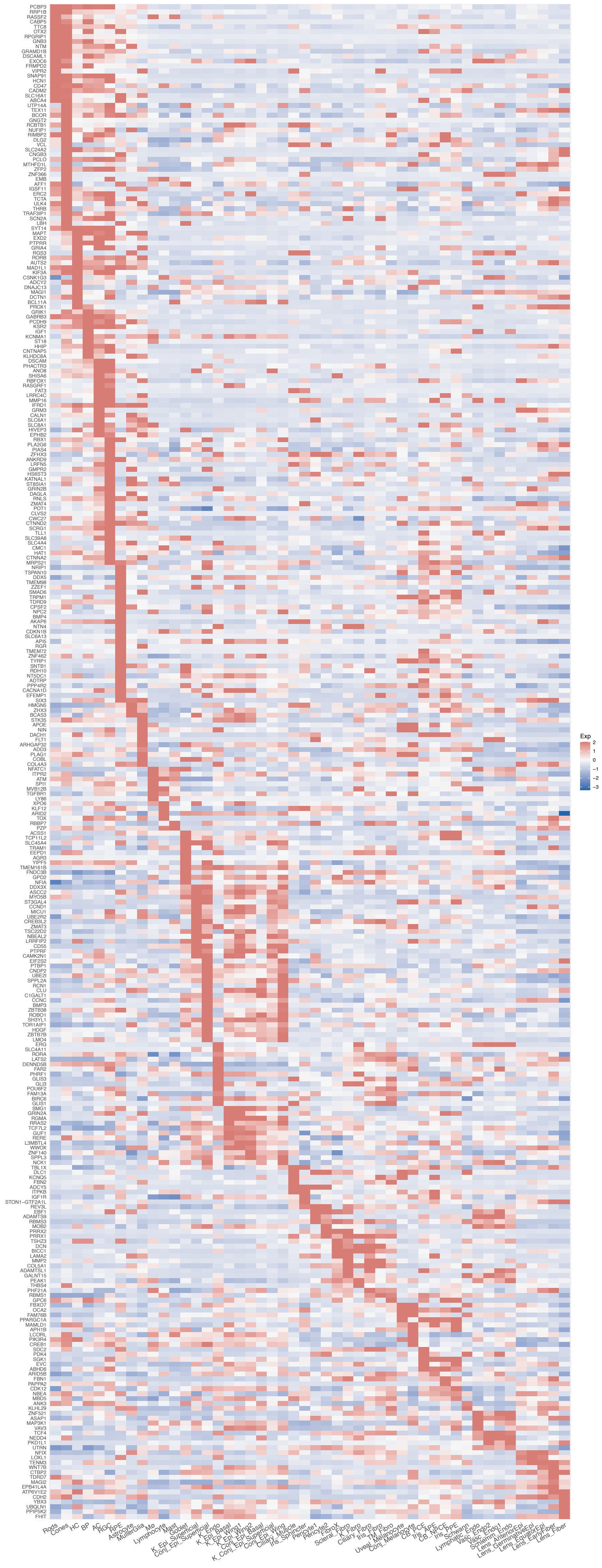

### FIGURE S6E - Primary Open Angle Glaucoma (POAG)

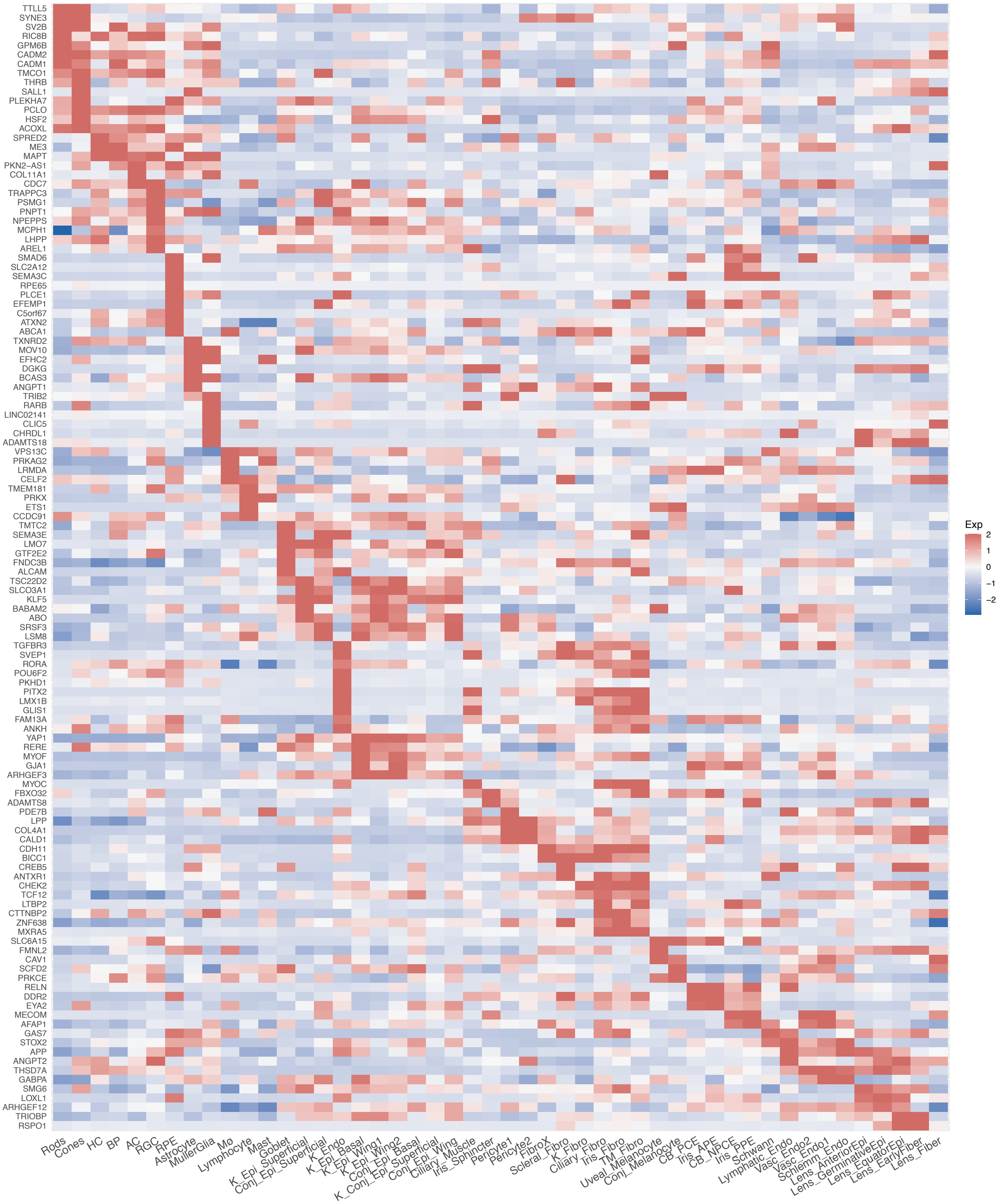

### FIGURE S6F - Refractive Error

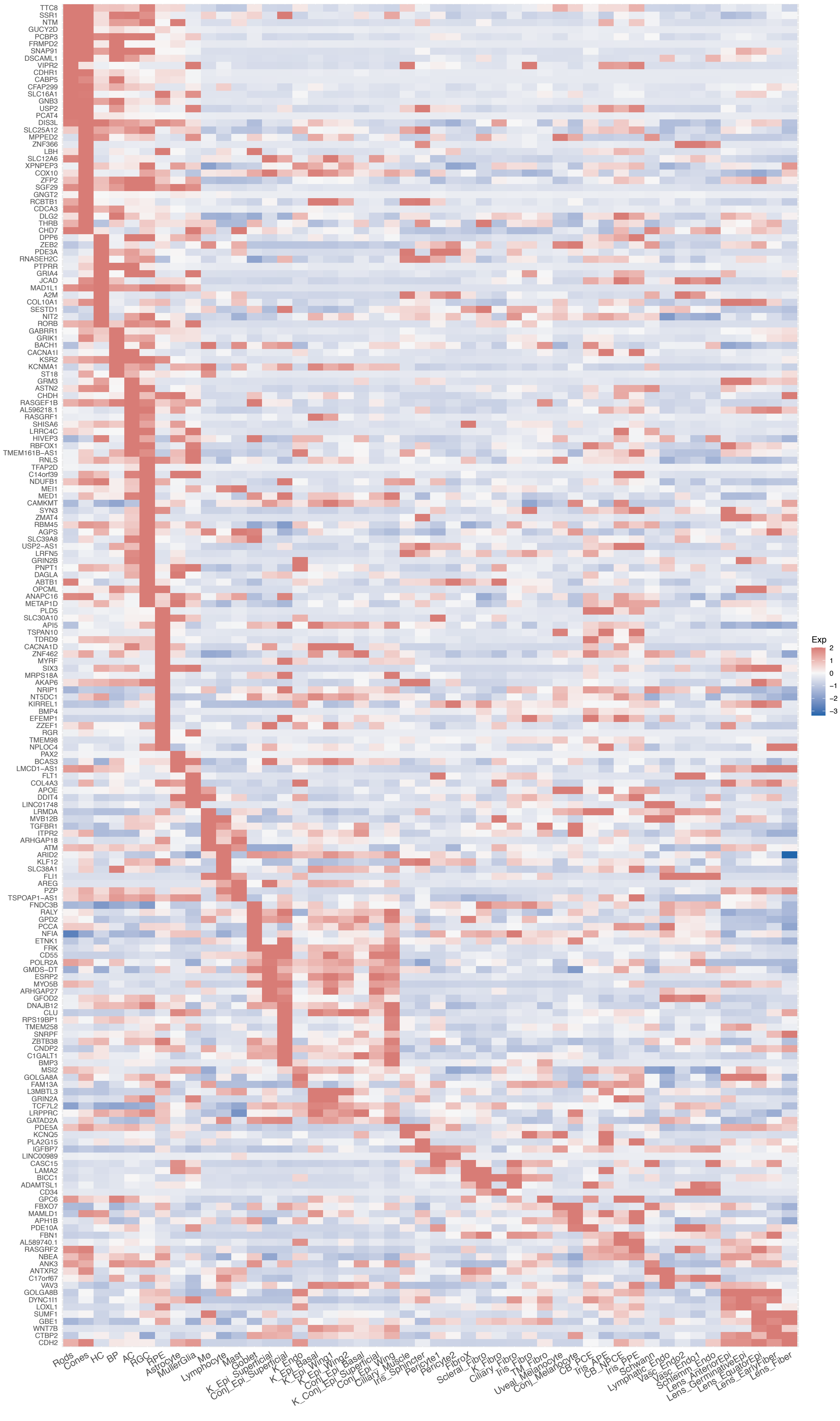

### FIGURE S6G - Vertical Cup to Disc Ratio (VCDR)

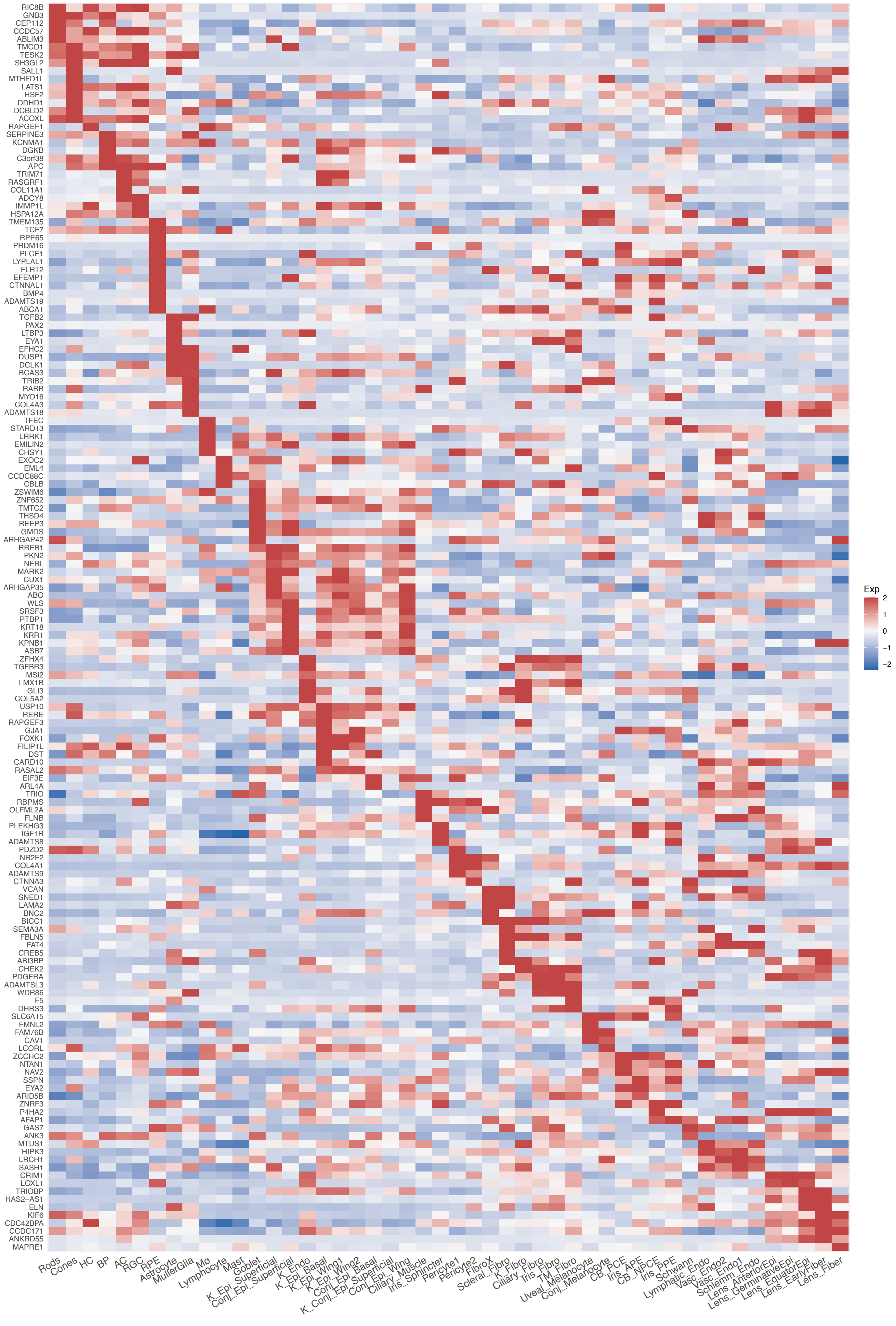

### Figure S6H - Cataract

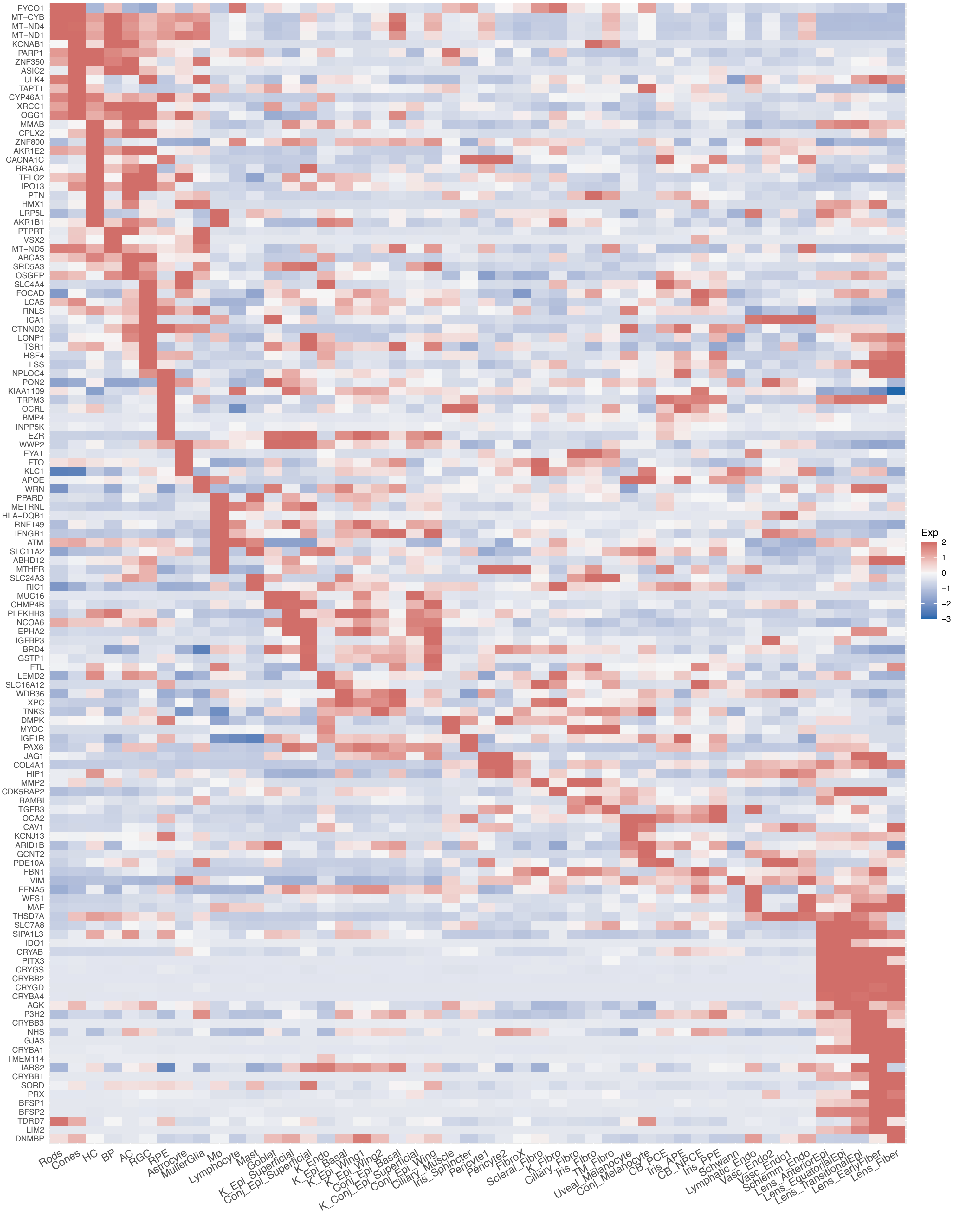

#### Supplementary Table 1. Donor Information (RNAseq)

| Label | Source | Age | Sex | Cause of Death | DTP (hours) | Tissues processed for RNAseq (10X version chemistry) |
|---|---|---|---|---|---|---|
| Pt2 Rt | MGH | 78 | M | Metastatic melanoma to brain | 14 | CB(V3) |
| Pt14-Rt | MGH | 33 | M | Colorectal cancer | 4 | Lens(V3.1) |
| 0216-20 OD | UT | 66 | F | Sepsis, end stage liver disease | 4.5 | CB(V3)<br>Iris(V3.1) |
| 0216-20 OS | UT | 66 | F | Sepsis, end stage liver disease | 4.5 | Cornea(V3), Iris(V3.1), CSW(V3.1), TM(V3.1)<br>Lens (V3.1) |
| 0218-20 OS | UT | 65 | M | Cardiogenic shock | 5.5 | TM(V3.1), Lens(V3.1), Cornea(V3), Iris(V3.1)<br>CSW(V3.1) |
| 0220-20 OS | UT | 47 | F | Metastatic brain cancer | 3.5 | TM(V3.1), Lens(V3.1), Cornea(V3), Iris(V3.1)<br>CSW(V3.1), CB(V3) |
| 0235-20 OS | UT | 30 | M | Respiratory failure | 5 | TM(V3.1), Lens(V3.1), Cornea(V3), Iris(V3.1)<br>CSW(V3.1), CB(V3) |
| 0822-19 | UT | 41 | M | Acute Cardiac Event | 5.3 | Macula (V3.1) |

DTP = Death to Processing Time

**Supplementary Table 2. Donor Information (Histology)**

| Label | Source | Age | Sex | COD | DTP (hr) | RNA *in situ* | IHC |
|---|---|---|---|---|---|---|---|
| 1041-16 | Utah | 43 | F | Sepsis | 5.5 | N/A | PDPN, PECAM1, DES, AQP1 |
| 1094-20 | Utah | 58 | M | Pancreatic cancer | 3.5 | ANGPTL7, BCAS1, LGR6, MECOM, PDGFC, NTRK2 | AQP1, RELN, LRP2 |
| Pt9-IrCB | MGH | 53 | F | Interstitial Lung Disease | 5 | N/A | CRB1, PDPN, PECAM1 |
| 1093-20 | Utah | 66 | F | Cardiac arrest | 4.5 | MECOM, LGR6, KRT12, RARRES1, CACNA1A, ATP8B4, BCAS1, NECTIN4, TOP2A, LAMA3, UCHL1, SLC1A2, GRIA4, ETNPPL, GPR160, CAV1 | MLANA, MUC5AC |
| Pt7-K | MGH | 52 | F | Brain hemorrhage; metastatic brain cancer | 4 | N/A | KRT78 |
| HCS13 | Lions | 22 | F | Cystic Fibrosis | 10 | N/A | MLANA |
| 0414-21 | Utah | 30 | M | Cardiac Arrest | 4.5 | BMP5, C7, PI16, NEB, PPP1R1B, TMEFF2, UNC5D, NELL2, PKHD1L1, FN1 | N/A |
| 0245-21 | Utah | 60 | F | Respiratory Arrest | 4.5 | NOTCH3, ID4, NDUFA4L2, ADCY3, WFDC2, ENTPD1, BMP5, ETNPPL, PAX3, MET, KIT, LEF1, FN1, GRIA4, ANGPTL7, PPP1R1B | N/A |

DTP = Death to Processing Time

**Supplementary Table 3. Antibody & RNA *in situ* Probe Information**

| **Antibody/Probe** | **Supplier** | **Catalog #** |
|---|---|---|
| Sheep Polyclonal anti-Podoplanin | R&D Systems | AF3670 |
| Rabbit Polyclonal anti-CD31/PECAM-1 | Novus Biologicals | NB100-2284 |
| Goat Polyclonal anti-Desmin | R&D Systems | AF3844 |
| Rabbit Polyclonal anti-Aquaporin-1 | Proteintech | 20333-1-AP |
| Rabbit Polyclonal anti-LRP2 | Lifespan Biosciences | LS-B15008 |
| Mouse Monoclonal anti-Reelin | Developmental Studies Hybridoma Bank | TB5 |
| Rabbit Polyclonal anti-CRB1 | Bioss Antibodies | BS-14045R |
| Mouse Monoclonal anti-Melan-A | Proteintech | 60348-1-lg |
| Mouse Monoclonal anti-Mucin 5AC | Abcam | ab3649 |
| Rabbit Polyclonal anti-Keratin 78 | Novus Biologicals | NBP1-93671 |
| Rabbit Polyclonal anti-CA3 | Lifespan Biosciences | LS-B9781 |
| RNAscope® Probe-Hs-ANGPTL7 | Advanced Cell Diagnostics | 552811 |
| RNAscope® Probe-Hs-BCAS1 | Advanced Cell Diagnostics | 525781-C3 |
| RNAscope® Probe-Hs-LGR6 | Advanced Cell Diagnostics | 410461-C2 |
| RNAscope® Probe-Hs-MECOM | Advanced Cell Diagnostics | 518021 |
| RNAscope® Probe-Hs-PDGFC | Advanced Cell Diagnostics | 404281 |
| RNAscope® Probe-Hs-NTRK2 | Advanced Cell Diagnostics | 402621-C2 |
| RNAscope® Probe-Hs-KRT12 | Advanced Cell Diagnostics | 506311-C2 |
| RNAscope® Probe-Hs-RARRES1 | Advanced Cell Diagnostics | 494851 |
| RNAscope® Probe-Hs-CACNA1A | Advanced Cell Diagnostics | 558581 |
| RNAscope® Probe-Hs-ATP8B4 | Advanced Cell Diagnostics | Custom[1] |
| RNAscope® Probe-Hs-NECTIN4 | Advanced Cell Diagnostics | 562061 |
| RNAscope® Probe-Hs-TOP2A | Advanced Cell Diagnostics | 470321 |
| RNAscope® Probe-Hs-LAMA3 | Advanced Cell Diagnostics | 530681 |
| RNAscope® Probe-Hs-UCHL1 | Advanced Cell Diagnostics | 594281 |
| RNAscope® Probe-Hs-SLC1A2 | Advanced Cell Diagnostics | 444721-C3 |

| | | |
|---|---|---|
| RNAscope® Probe-Hs-GRIA4 | Advanced Cell Diagnostics | 464671 |
| RNAscope® Probe-Hs-ETNPPL | Advanced Cell Diagnostics | 492121-C3 |
| RNAscope® Probe-Hs-GPR160 | Advanced Cell Diagnostics | 482681 |
| RNAscope® Probe-Hs-CAV1 | Advanced Cell Diagnostics | 452071-C2 |
| RNAscope® Probe-Hs-BMP5 | Advanced Cell Diagnostics | 472461 |
| RNAscope® Probe-Hs-C7 | Advanced Cell Diagnostics | 534791-C3 |
| RNAscope® Probe-Hs-PI16 | Advanced Cell Diagnostics | 569181 |
| RNAscope® Probe-Hs-NEB | Advanced Cell Diagnostics | 554391-C3 |
| RNAscope® Probe-Hs-PPP1R1B | Advanced Cell Diagnostics | 477021-C2 |
| RNAscope® Probe-Hs-TMEFF2 | Advanced Cell Diagnostics | 519741 |
| RNAscope® Probe-Hs-UNC5D | Advanced Cell Diagnostics | 459991 |
| RNAscope® Probe-Hs-NELL2 | Advanced Cell Diagnostics | Custom[2] |
| RNAscope® Probe-Hs-PKHD1L1 | Advanced Cell Diagnostics | Custom[3] |
| RNAscope® Probe-Hs-BTNL9 | Advanced Cell Diagnostics | 430351 |
| RNAscope® Probe-Hs-FN1 | Advanced Cell Diagnostics | 310311-C2 |
| RNAscope® Probe-Hs-NOTCH3 | Advanced Cell Diagnostics | 558991-C2 |
| RNAscope® Probe-Hs-ID4 | Advanced Cell Diagnostics | 466371-C3 |
| RNAscope® Probe-Hs-NDUFA4L2 | Advanced Cell Diagnostics | 567011-C3 |
| RNAscope® Probe-Hs-ADCY3 | Advanced Cell Diagnostics | 441671 |
| RNAscope® Probe-Hs-WFDC2 | Advanced Cell Diagnostics | 524781 |
| RNAscope® Probe-Hs-ENTPD1 | Advanced Cell Diagnostics | 474181-C2 |
| RNAscope® Probe-Hs-PAX3 | Advanced Cell Diagnostics | 562711-C2 |
| RNAscope® Probe-Hs-MET | Advanced Cell Diagnostics | 431021-C3 |
| RNAscope® Probe-Hs-KIT | Advanced Cell Diagnostics | 606401-C3 |
| RNAscope® Probe-Hs-LEF1 | Advanced Cell Diagnostics | 412991 |

[1] 20 ZZ probe targeting 1119-2158 of NM_024837.4
[2] 20 ZZ probe targeting 668-1660 bp of NM_001145108.2
[3] 20 ZZ probe targeting 4456-5459 bp of NM_177531.6
